# Long-term homogenization of vascular plant and lichen communities across Fennoscandian heathlands and tundra is connected to the expansion of an allelopathic dwarf shrub

**DOI:** 10.1101/2024.12.27.630519

**Authors:** Tuija Maliniemi, Petteri Kiilunen, Kari Anne Bråthen, Jutta Kapfer, Torunn Bockelie Rosendal, John-Arvid Grytnes, Patrick Saccone, Risto Virtanen

## Abstract

Boreal and tundra plant communities are expected to change in biodiversity due to increasing global change pressures such as climate warming. One long-term scenario is increasing compositional similarity, i.e., biotic homogenization, which has been relatively little studied in high-latitude plant communities. Here, we study how the composition and diversity of heathland and tundra plant communities have changed in northern Fennoscandia over several decades. In 2013–2023, we resurveyed 275 historic vegetation plots, originally surveyed in 1964–1975, with percentage covers for vascular plant, bryophyte and lichen species. We analyzed temporal changes in community composition and diversity across the study area and in different habitat types, biogeographic zones, and along the continentality-humidity gradient. We found a strong trend across the study area, with plant communities becoming more similar in composition over the decades. The observed homogenization was associated with compositional changes in vascular plant and lichen communities, and in particular with the encroachment of the evergreen dwarf shrub *Empetrum nigrum*. In comparison to vascular plants and lichens, the diversity of bryophytes generally remained more stable over time. Our findings suggest that Fennoscandian heathland and tundra vegetation is transforming towards a more homogenous evergreen dwarf shrub dominated system, which may threaten ecosystem multifunctionality. Our results highlight the importance of exploring biodiversity among different metrics and growth forms to understand the overall changes in heathland and tundra biodiversity.

## Introduction

Biodiversity change is expected to have a considerable impact on the structure and functions of high-latitude biomes, where rapidly changing climate (Rantanen et al. 2022) is a key driver reshaping biodiversity (Sala et al. 2000, Niittynen et al. 2018, García Criado et al. 2024). While changes in alpha-diversity (e.g. species richness) have been the main focus across spatial and temporal scales in the tundra (e.g. Wilson & Nilsson 2009, Vellend et al. 2017), long-term changes in compositional similarity, or beta-diversity, have gained less attention until lately (Bråthen et al. 2024, García Criado et al. 2024). One scenario for beta-diversity is a decline in compositional dissimilarity, i.e., biotic homogenization, in which communities become more similar regarding taxonomy, functional roles or genetic variation (McKinney & Lockwood 1999, Olden & Rooney 2006). This, in turn, is expected to have a negative impact on ecosystem resilience (Olden et al. 2004) and the provision of multiple ecosystem functions, i.e. multifunctionality (Hautier et al. 2018, Mori et al. 2018, Liu et al. 2021). Biotic homogenization can result from unique species disappearing (Olden & Rooney 2006) or common species invading and becoming regionally dominant (Olden & Rooney 2006, McCune & Vellend 2012, Socolar et al. 2016). Alternatively, changing local conditions and environmental gradients can increase among-site dissimilarity, leading to compositional differentiation, i.e., biotic heterogenization. Importantly, changes in beta-diversity can be decoupled from changes in alpha-diversity (Li et al. 2020, Dornelas et al. 2023) and regional homogenization can occur even as local diversity increases (e.g., McCune & Vellend 2012, Finderup Nielsen et al. 2019). Increasing knowledge of long-term beta-diversity change will broaden the understanding of biodiversity change in high-latitude environments and the potential consequences for the ecosystem multifunctionality. In addition, it will contribute to estimating ecosystem state and condition (e.g. Pedersen et al. 2021) and reaching the targets of national and international biodiversity programmes and strategies (IPBES 2019, CBD 2022).

Compositional dissimilarity among plant communities is essentially maintained by environmental and landscape heterogeneity, which supports the provision of ecosystem functions and services across space and time (Isbell et al. 2011, Hautier et al. 2017, Mori et al. 2018). In boreal-Arctic environments, homogenization may result from various drivers that alter environmental conditions, such as changing climate (Walker et al. 2006, Niittynen et al. 2020), or from biotic interactions, such as herbivore densities (Rooney 2009, Ravolainen et al. 2010) or plant competition (Bråthen et al. 2024). When such drivers operate on a large scale, they can decrease the variation of local conditions; this often benefits generalist species across localities and decreases the local dominance of specialist species, which reduces beta diversity (Hillebrand et al. 2008). Thus, the decreased spatial variation among species communities and changes towards regional dominance of a few species likely decrease multifunctionality across the landscape.

Shrub expansion is one of the best documented and ecologically important outcomes of global climate warming in the tundra (Elmendorf et al. 2012, Myers-Smith et al. 2020). It has been predicted to homogenize tundra plant communities by fostering the dominance of already abundant shrub species, which may pose far-reaching impacts to the ecosystem structure and functions (Mod et al. 2016, Bråthen, Gonzalez & Yoccoz, 2018, Steward et al. 2018). Even though a recent meta-analysis indicated no homogenization of vascular plant communities across the Arctic (García Criado et al. 2024), there is limited knowledge of such trends at regional and local scales. The observed large-scale shrub expansion often refers to the increase in tall, deciduous shrubs, but it is also linked to the increase in evergreen shrubs (Elmendorf et al. 2012, Vowles et al. 2019), such as the dwarf shrub *Empetrum nigrum* (Wilson & Nilsson 2009, Vowles et al. 2017, Maliniemi et al. 2018, Stark et al. 2021, Bråthen et al. 2024, Tuomi et al. 2024). *E. nigrum* has encroached vigorously in parts of the circumpolar Arctic and has established dominance in many habitat types (Maliniemi et al. 2018, Bråthen et al. 2024, Maliniemi et al. 2024). *E. nigrum* has a number of traits that make it a strong competitor, with several negative impacts on plant community structure and diversity, as well as ecosystem functions and properties (Tybirk et al. 2000, Tuomi et al. 2024). The allelopathic effect of *E. nigrum* inhibits the establishment of other species (Bråthen et al. 2010, González et al. 2015), even extending to neighbouring habitats (Pilsbacher et al. 2021), and the phenolic content of its leaves and litter further slows decomposition and soil nutrient cycling (Tybirk et al. 2000). In addition, dense, mat-forming clonal growth makes *E. nigrum* a superior competitor, preventing other species to establish (Tybirk et al. 2000). Increasing dominance of *E. nigrum* poses a threat to plant diversity in the tundra, where it has already been reported to reduce species diversity (Olofsson et al. 2009, Wilson & Nilsson 2009, Pellissier et al. 2010, Bråthen & Ravolainen 2015), habitat type diversity (Bråthen et al. 2024), and to deteriorate pasture quality (Tuomi et al. 2024). However, we still lack evidence on how the expansion of *E. nigrum* is reflected in the compositional dissimilarity of tundra plant communities in the long-term.

While diversity changes in vascular plants have received most of the attention (e.g. Garcia Criado et al. 2024), long-term diversity changes in non-vascular cryptogams, i.e., bryophytes and lichens, have received less attention. This gap in knowledge is considerable given that these species form a substantial part of tundra biodiversity, and provide a variety of ecosystem functions related to, for example, nutrient cycling, soil and biomass production, nitrogen fixation and herbivory diet (Cornelissen et al. 2007, Turetsky et al. 2012, Asplund & Wardle 2017, Lett et al. 2021). Although certain responses of cryptogams are generally well studied, such as the negative response of lichen cover and biomass to reindeer trampling and foraging (Kumpula et al. 2014, Bernes et al. 2015), it is still poorly understood how the composition of bryophyte and lichen communities has changed over long time periods and to what extent these different growth forms are similar in their long-term biodiversity patterns.

In this study, we analyze long-term changes in the diversity of Fennoscandian oligotrophic treeless heathland and tundra plant communities and whether they are dependent on different environmental conditions. We use vegetation resurvey data that include abundance information for vascular plant, bryophyte and lichen species and cover spatially extensive area and temporally over five decades. We ask, 1) has beta-diversity (compositional dissimilarity) declined over time across the study system, considering all species together as well as different growth forms (vascular plants, bryophytes and lichens) separately, 2) are the changes comparable within different environmental conditions, i.e. biogeographic zones, degree of continentality-humidity and habitat types and 3) are changes in beta-diversity linked to the changes in species richness, evenness or abundance of dominant species? We hypothesize that Fennoscandian tundra communities have become compositionally more similar over time, but these responses may depend on environmental variation and may not be parallel among different biodiversity metrics or growth forms.

## Materials and Methods

### Study area and the resurvey of plant communities

We resurveyed a historical vegetation sample plot data from treeless mountain heath and tundra highlands in northern Fennoscandia, spanning a large gradient along latitude and longitude (Fig. 1). The original vegetation survey included 987 plots and was conducted during 1964–75 by Matti Haapasaari (1988) to describe and classify oligotrophic mountain heath and tundra vegetation of the area. This vegetation type covers extensive areas in northern Fennoscandia and is dominated by dwarf shrubs (ericoids) and low shrubs (*Betula nana*) often with a considerable share of bryophytes and lichens, which frequently outnumber vascular plants (Haapasaari 1988, Virtanen et al. 2016, Maliniemi et al. 2018). Semi-domesticated reindeer is the main large herbivore in the area, where their numbers have been relatively high before and during the whole study period (Bernes et al. 2015, Maliniemi et al. 2018, Stark et al. 2023), while small rodents (e.g., voles and lemmings) are important smaller herbivores in the area (Soininen et al. 2024). Study sites include site types from northern boreal to oroarctic, elevation ranging from 0 to 720 m.a.s.l., and climatic conditions vary from oceanic to more continental (Haapasaari 1988, Rantanen et al. 2023). Since 1950, the study area has experienced an approximate 0.15-0.30°C increase in annual mean temperature per decade and an approximate 1-4 day increase in thermal growing season length per decade (Rantanen et al. 2023).

**Figure 1.**
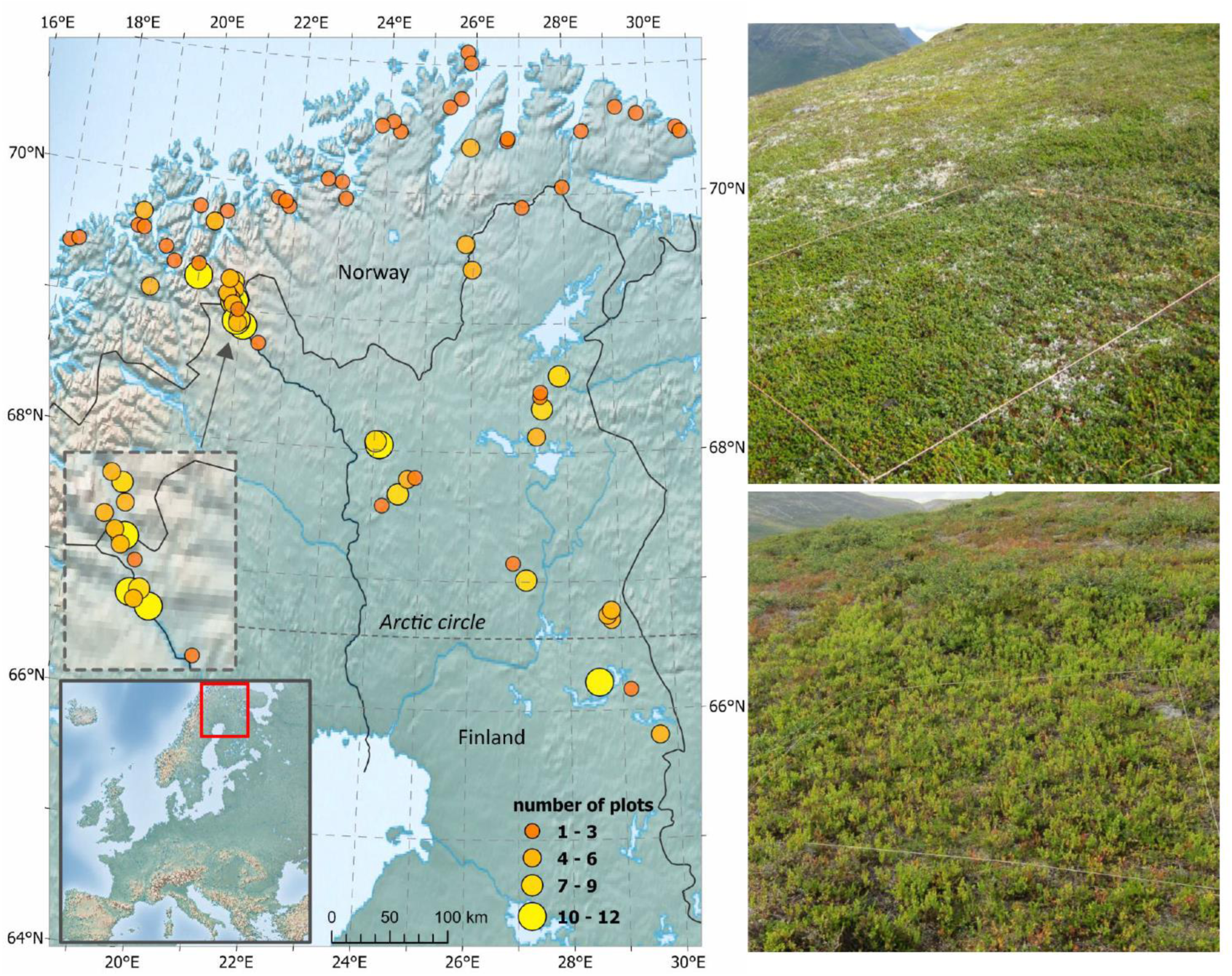
The location of the 275 sample plots in Northern Fennoscandia. Each point on the map includes one to twelve independent vegetation sample plots. Photos illustrate the two most common habitat types of oligotrophic heathlands in the area, *Empetrum nigrum*-dominated (upper right) type and *Vaccinium myrtillus*- dominated (lower right) type.

In 2013–2023, we resampled a total of 275 vegetation plots using the same methods as in the original survey. The relocation of the plots was based on detailed plot-specific location information (the description of the area and the surroundings, elevation, slope, aspect and the extent of the specific site type) in the original publication (Haapasaari, 1988). Thus, plots can be considered as quasi-permanent (Kapfer et al. 2017). Plots with uncertain relocation information were not included. None of the plots included in this study showed signs of direct human disturbance. Similar to the original survey, plots of 2 x 2 m were placed within a specific habitat type, often extending over a large area, and on spots representative of species composition of the habitat type. Vascular plant, bryophyte, and lichen species were recorded from each plot using a percentage cover scale (0.1, 0.25, 0.5, 1, 2, 3, 5, 10, 20,…90, 100%). Each species was given an absolute cover estimate, so the total cover of all species in a plot can exceed 100%. The relocation error is inherent in the resurveys of quasi-permanent plots, but it was minimized according to Kapfer et al. (2017) to guarantee qualified comparisons between the original and resurveyed data. This included, e.g., resampling phenologically similar time to the original survey and using only well-trained field botanists to minimize observer error. Moreover, we did not make pair-wise comparisons of plots but studied habitat-level changes for which quasi-permanent plot data is well suited (Alfonsi et al. 2017). Before the analysis, species data was harmonized between the surveys. To reduce possible errors related to changed taxonomy or misidentification due to high number of small, infrequent species, moss and lichen species were treated at the genus level, except for the most common and abundant, so-called “reindeer lichens” (Table S1). Liverworts were treated together (*Hepaticae*), except for the large-sized and frequent species *Barbilophozia lycopodioides* and *Ptilidium ciliare*. Crustaceous lichens growing on the ground were given a common total cover value. Lichens growing on rocks were not considered. If bryophyte or lichen taxa included only one species (e.g., *Pleurozium schreberi*) the species-specific name is used. The total number of resulting taxa is 173 of which 118 are vascular plants, 35 bryophytes and 20 lichens (Table S2). It should be noted that the real number of non-vascular cryptogams is much higher due to the merged species (Table S1).

To test if diversity changes were spatially limited to certain environmental conditions, we classified sample plots by their biogeographical distribution, based on a classification made by Haapasaari (1988, Fig. S1). Biogeographical classes include northern boreal (plot n = 96), hemioroarctic (n = 84) and oroarctic site types (n = 95) (class names updated using Virtanen et al. 2016). Northern boreal types are characteristic of low-elevation heathlands throughout the study area, transitioning to hemioroarctic and oroarctic types as elevation increases. Plots were also classified based on the degree of continentality-humidity (Haapasaari, 1988, Fig. S1). Continentality‒humidity classes include subcontinental/arid (n = 47), indifferent (transition between subcontinental and oceanic, n = 111) and oceanic/humid communities (n = 117). It should be noted that this classification is not a direct measure of distance to the ocean but includes local humidity: these classes are characterised by either substantial lichen cover (subcontinental), bryophyte cover (oceanic) or both equally present (indifferent). Importantly, classifications are independent of each other, for example, oroarctic communities are found from subcontinental to oceanic conditions (Table S3, Fig. S1). Finally, plots were classified by their habitat types (Haapasaari 1988) that are determined by mesotopography and resulting snow cover and soil moisture gradients (Haapasaari, 1988; Maliniemi et al. 2018; Kuusisto et al. 2024). Of the plots, 104 were originally of *Empetrum nigrum*-dominated types, 87 *Vaccinium myrtillus*-types, 31 *Betula nana*-types, 29 *Calluna vulgaris*-type and 24 relatively barren, windswept types. Based on IUCN classification, first three types are considered near threatened and two latter vulnerable in Finland (Kontula & Raunio, 2019). *E. nigrum*- and windswept types generally represent chionophobus heath types, i.e. thriving under less snow, while *V. myrtillus-*, *B. nana-* and *C. vulgaris*-types represent chionophilous types that require more snow (Haapasaari, 1988). The original habitat type was typically either fully or partly recognizable in the resurvey and the plot was placed on an average spot in terms of community composition, as in the original survey. The uneven number of resurveyed plots within the habitat types reflects their commonness and thus represents the true proportion of types in the oligotrophic Fennoscandian tundra: *E. nigrum*- and *V. myrtillus*-types are the most common types, while *B. nana*-type is typical of northern and higher elevation sites, *C. vulgaris*-type is typical of southern, coastal and lower elevation sites and small patches of windswept sites are the least common (Haapasaari, 1988, Fig. S1). The small number of the three latter may add some uncertainty to the observed changes in these habitats, but strong trends can be taken as indicative of changes that have occurred over time.

### Statistical analyses

To study whether communities had become more similar (homogenized) or more dissimilar (heterogenized) over time, we analyzed the multivariate homogeneity of group dispersions (PERMDISP2, Anderson et al. 2006). Species abundances (% cover) and Bray-Curtis dissimilarities were used as a distance structure in all the analyses. We used a function betadisper from the R package *vegan* (Oksanen et al. 2024) to calculate the average distance of group members (plots in the original survey or the resurvey) to their group centroid (original survey or resurvey) in a multivariate space. First, we calculated group centroid (type ‘spatial median’ in the function) for old and new communities in a multivariate space using all plots and all taxa. Then, using permutation test (with 9999 permutations), we tested whether the group dispersions were homogenous over time, i.e., if there were difference in the average distance to the spatial median between the surveys. Compositional shifts were visualized with PCoA ordination plots. To test if the cover (%) of most dominant species or different growth forms correlated with compositional shifts, we added correlation vectors to the PCoA for the cover of *E. nigrum*, *V. myrtillus*, *V. vitis-idaea*, *B. nana*, *C. vulgaris* as well as for the total cover of vascular plants, bryophytes and lichens. The goodness-of-fit of vectors was assessed with 9999 permutations. Second, to test if the observed changes were linked to certain environmental conditions, we calculated spatial medians for the subsets of data, i.e., for different biogeographical zones, continentality classes and habitat types, and tested with PERMDISP2 whether average distance to spatial median had changed over time within the different zones, classes and habitat types. Even though the subsets were tested separately, they were visualised together to compare the direction of compositional change. Finally, to explore if the observed patterns in compositional changes were driven by certain species groups, all the analyses described above were made separately for vascular plant, bryophyte and lichen communities.

To study changes in alpha diversity, we explored its two components—species richness and evenness (dominance)—as they may be linked to the mechanism through which homogenization or heterogenization occurs. For example, homogenization accompanied by decreasing species richness relates to the disappearance of unique species, while simultaneous changes in evenness indicate changes in either local or regional dominance. Species richness and evenness were calculated across the study area and for each biogeographical zone, continentality class and habitat type during both surveys. We calculated richness for all taxonomic groups together and separately for vascular plants, bryophytes and lichens. Paired t-tests with 9999 permutations were used to test changes over time, using the package *RVAideMemoire* (Herve 2023). To keep it conservative due to multiple testing, we used a p-value threshold of 0.01 when interpreting the results.

Finally, to estimate species-specific contributions to compositional changes, we identified those species that had changed the most over time across all sites. For this, we used paired t-tests between the original and resurveyed species abundances. T-tests were performed with 9999 permutations to allow testing of species abundances with non-normal distributions, and to further correct p-values for multiple testing (Legendre, 2019). Multiple testing was taken into account in two conservative ways. First, we set the significance level after permutations to 0.01 (instead of commonly used 0.05) and then used Holm correction for multiple testing. Both permuted and corrected p-values are displayed in the results. Analyses were made using the package *adespatial* (Dray et al. 2024).

## Results

When all sites and all taxa (vascular plants, bryophytes and lichens) were considered, the average distance of plant communities to the group centroid (i.e., spatial median) in a multivariate space decreased over time, indicating a homogenization of tundra plant communities across the study area (Fig. 2a, Fig. 3, Table S4). Homogenization occurred within all growth forms (Fig. 3). In general, plant communities showed a temporal shift towards increased *E. nigrum* coverage as well as a shift away from higher lichen coverage in the original survey (Fig. 2a).

**Figure 2.**
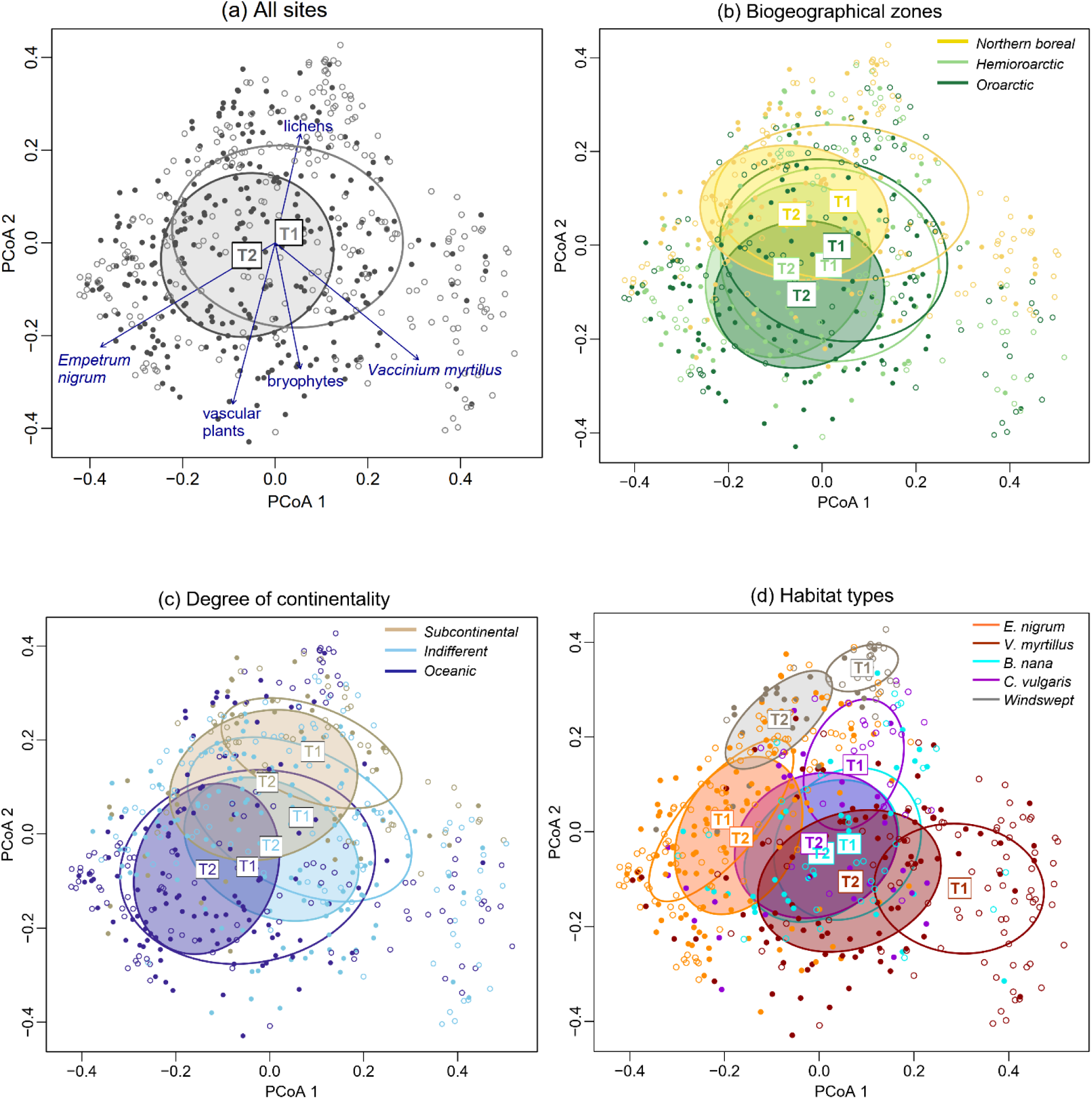
Principal coordinate analysis (PCoA) based on Bray-Curtis dissimilarities and the abundances (cover %) of all species (vascular plants, bryophytes and lichens) on vegetation plots (points). In each panel, labels point the location of spatial median of plots, surrounded by 1 standard deviation ellipses, during the original survey (T1, hollow symbols) and resurvey (T2, filled symbols). Spatial median is calculated for both surveys **a)** across all study sites (with statistically significant correlation vectors for the cover of *E. nigrum*, r^2^ = 0.87, p = 0.001; *V. myrtillus*, r^2^ = 0.72, p = 0.001; vascular plants, r^2^ = 0.58, p = 0.001; bryophytes, r^2^ = 0.35, p = 0.001 and lichens, r^2^ = 0.26, p = 0.001; see Table S5 for non-significant vectors), **b)** in different biogeographical zones, **c)** in different continentality classes and **d)** in different habitat types. Note that panels are directly comparable, as they are based on the same PCoA.

**Figure 3.**
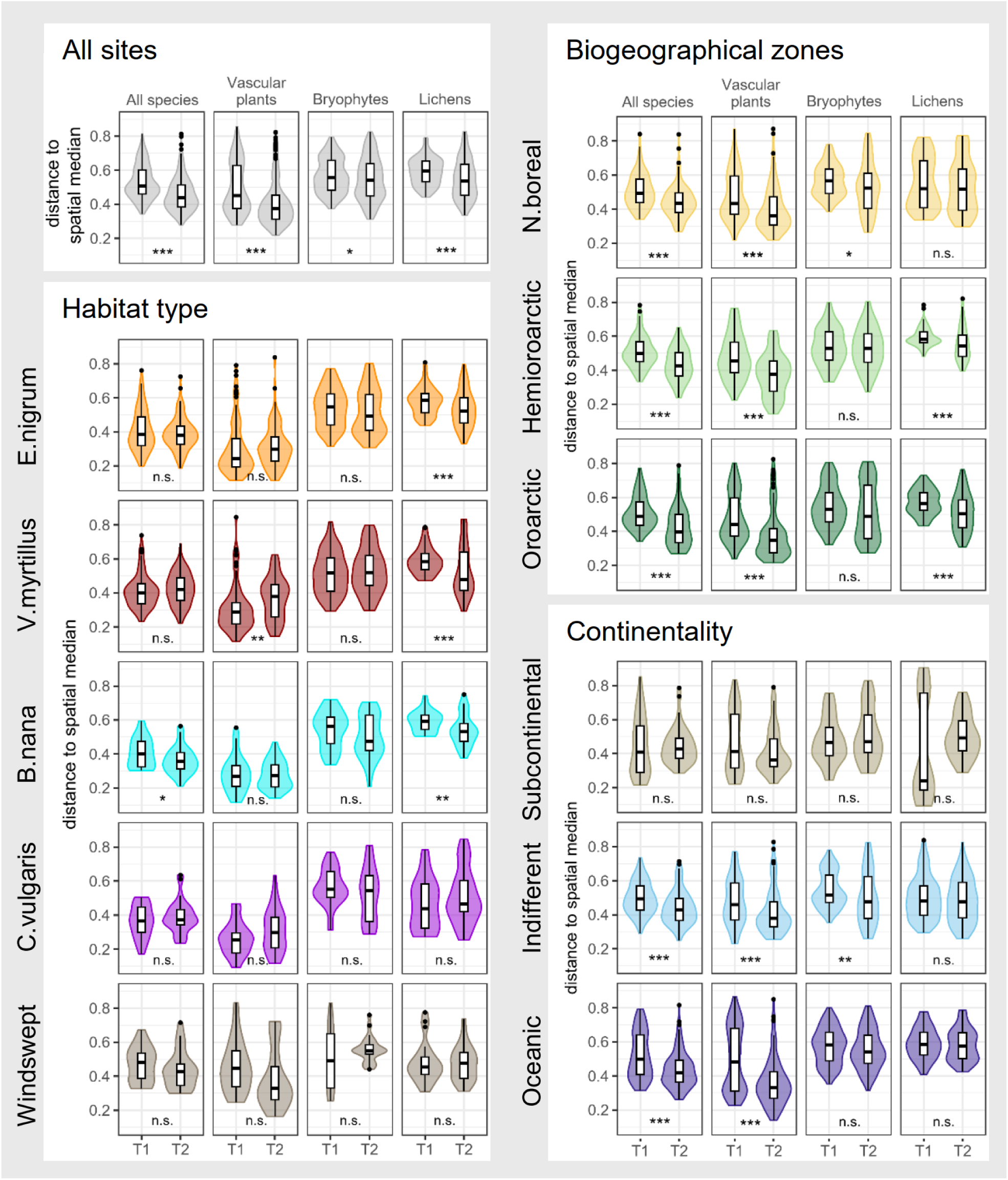
Compositional dissimilarity across all sites and within biogeographical zones, continentality classes and habitat types during the original survey in 1960s (T1) and the resurvey in 2010s (T2). Dissimilarity is measured as a distance to spatial median in a multivariate space for the whole community composition and separately for vascular plants, bryophytes and lichens. The effect of time on the average distance to spatial median is indicated as *p < 0.05, **p < 0.01, ***p < 0.001, n.s. = not significant.

A homogenization pattern was found within all biogeographical zones (Fig. 2b, Fig.3) when all taxa were treated together. Among growth forms, vascular plant communities homogenized within each zone, bryophyte communities in northern boreal zone and lichen communities in hemioroarctic and oroarctic zones. Of continentality classes, subcontinental communities showed no homogenization over time, neither across all taxa or within any growth form (Fig. 3). Indifferent and oceanic communities showed homogenization across all taxa and within vascular plant communities. Also, bryophyte communities homogenized in indifferent conditions.

Of the habitat types, only *B. nana*-dominated communities experienced within-habitat homogenization, when all taxa were treated together (Fig. 3). However, there were some growth form-specific responses. Vascular plant communities heterogenized in *V. myrtillus-*types and lichen communities homogenized in *E. nigrum*, *V. myrtillus* and *B. nana-*types over time.

The richness of lichen taxa declined in northern boreal zone, in indifferent sites and in *C. vulgaris*-type (Table 1), but the magnitude of the decrease was not pronounced (Table S6). Evenness, in turn, increased across all sites and in *E. nigrum*-type, where it increased among vascular plants and lichens (Table 1). In addition, vascular plant evenness increased in northern boreal zone, in indifferent sites and in *V. myrtillus*-type, and lichen evenness in hemioroarctic and oroarctic sites and in subcontinental and indifferent sites. A decrease in evenness was only found for vascular plants in the windswept type. Bryophyte taxa showed no changes in richness or evenness over time (Table 1, Table S6). There were no major species-specific gains or losses over time, as those species gained or lost were typically recorded only a few times during either of the surveys (Table S2) (but see *Diphasiastrum complanatum*, Table 1).

**Table 1.** Summary of temporal dynamics in alpha diversity (species richness, SR; evenness, EVE) and beta diversity (compositional dissimilarity, DIS). Arrows indicate the direction of temporal change as follows: increase with p < 0.001 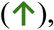 p < 0.01 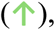 p < 0.05 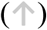 and decrease with p < 0.001 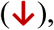 p < 0.01 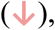 p < 0.05 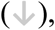 no change (-). See for values in Table S6 (alpha diversity) and in Fig. 3 and Table S4 (beta diversity).

|  | All sites |  |  | Habitat types |  |  |  |  |  |  |  |  |  |  |  |  |  |  |
| --- | --- | --- | --- | --- | --- | --- | --- | --- | --- | --- | --- | --- | --- | --- | --- | --- | --- | --- |
|  |  |  |  | E. nigrum |  |  | V. myrtillus |  |  | B. nana |  |  | C. vulgaris |  |  | Windswept |  |  |
|  | SR | EVE | DIS | SR | EVE | DIS | SR | EVE | DIS | SR | EVE | DIS | SR | EVE | DIS | SR | EVE | DIS |
| All species | - | ↑ | ↓ | - | ↑ | - | - | - | - | - | - | ↓ | ↓ | - | - | ↑ | - | - |
| Vascular plants | ↑ | ↑ | ↓ | - | ↑ | - | ↑ | ↑ | ↑ | - | - | - | - | ↑ | - | - | ↓ | - |
| Bryophytes | - | - | ↓ | - | - | - | - | - | - | - | - | - | - | - | - | - | - | - |
| Lichens | - | ↑ | ↓ | - | ↑ | ↓ | - | - | ↓ | - | - | ↓ | ↓ | - | - | - | - | - |
|  | Biogeographical zones |  |  |  |  |  |  |  |  | Continental-humidity |  |  |  |  |  |  |  |  |
|  | N. boreal |  |  | Hemioarctic |  |  | Oroarctic |  |  | Subcontinental |  |  | Indifferent |  |  | Oceanic |  |  |
|  | SR | EVE | DIS | SR | EVE | DIS | SR | EVE | DIS | SR | EVE | DIS | SR | EVE | DIS | SR | EVE | DIS |
| All species | - | - | ↓ | - | - | ↓ | - | - | ↓ | - | - | - | - | - | ↓ | - | ↑ | ↓ |
| Vascular plants | - | ↑ | ↓ | - | - | ↓ | - | - | ↓ | - | - | - | ↑ | ↑ | ↓ | - | - | ↓ |
| Bryophytes | - | - | ↓ | - | - | - | - | - | - | - | - | - | - | - | ↓ | - | - | - |
| Lichens | ↓ | - | - | - | ↑ | ↓ | - | ↑ | ↓ | - | ↑ | - | ↓ | ↑ | - | - | - | - |

Across all sites, the most pronounced changes in species mean cover (significant after correcting for multiple testing) were the increases in dwarf shrubs *B. nana*, *E. nigrum*, and *V. uliginosum*, increase of the tree *B. pubescens ssp. czerepanovii* and decrease of the dwarf shrub *V. myrtillus* (Table 2). Among the non-vascular cryptogams, the cover of bryophytes (*Hepaticae* and *Polytrichum* spp.) and lichens (*Stereocaulon* spp.), considered as pioneer species typical of open ground, decreased. The increase of *E. nigrum* was substantial across all biogeographical zones and continentality classes, with the largest increases occurring in the northern boreal zone and in subcontinental conditions (Table S7-8) and in *C. vulgaris*-, *V. myrtillus*- and windswept habitat types (Fig. 4). The decline of *V. myrtillus*, in turn, took place mainly in hemioroarctic and oroarctic zones, in oceanic conditions and in the *V. myrtillus-*type, in the latter of which it had become subordinate to *E. nigrum* (Fig. 4). The decrease of *Stereocaulon* spp. was profound in subcontinental conditions (Table S7-8) and in *E. nigrum-* and *V. myrtillus-*types (Fig. 4).

**Figure 4.**
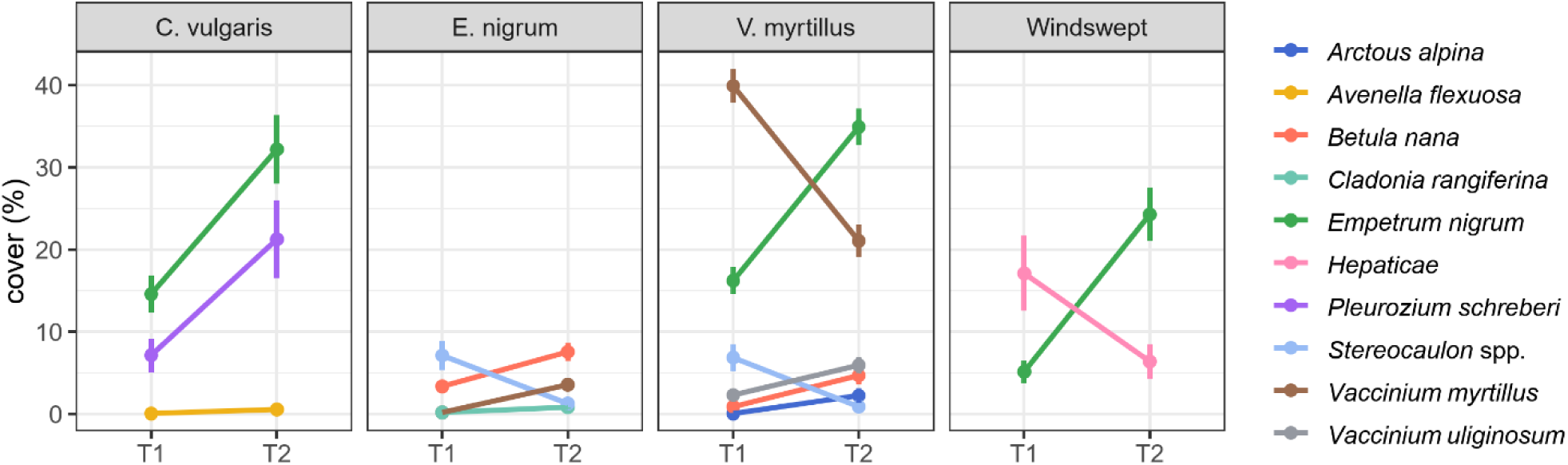
The most pronounced species-specific trends over time within different habitat types. Only species that showed significant change in mean cover after correcting for multiple testing are displayed (see Table S9 for all changes). For each species, mean cover and 95 % confidence intervals are given during the original survey (T1) and resurvey (T2). Of unchanged dominant species (not shown in the figure), *C. vulgaris* retained its position as a dominant species in *C. vulgaris* type over time (with a mean cover of 37 %) as did *E. nigrum* in *E.nigrum-*types (with a mean cover of 45 %).

**Table 2.** Species-specific changes across all sites over time. Species with significant cover changes (p.perm < 0.01) are listed. Significance is derived from permutational (9999) t-tests (p.perm) and corrected for multiple testing (p.corr). Species’ frequencies and mean covers in both original survey (T1) and resurvey (T2) and the observed change in cover (Δ) are given. All species are listed in Table S2. ‘+’ mean cover value is less than 0.1.

|  | All sites (n plots = 275, n species = 173) |  |  |  |  |  |
| --- | --- | --- | --- | --- | --- | --- |
| species | frequency T1 / T2 | cover (%) T1 / T2 | Δ cover | t.stat | p.perm | p.corr |
| Vascular plants |  |  |  |  |  |  |
| *Betula nana* | 157 / 160 | 6.7 / 9.8 | +3.1 | 4.4 | 0.001 | 0.017 |
| *Betula pubescens ssp. czerepanovii* | 4 / 30 | + / 0.3 | +0.3 | 2.3 | 0.001 | 0.017 |
| *Calamagrostis lapponica* | 65 / 43 | 0.3 / 0.1 | -0.1 | -1.8 | 0.010 | 1 |
| *Diphasiastrum complanatum* | 12 / 0 | 0.1 / 0 | -0.1 | -1.7 | 0.001 | 0.079 |
| *Empetrum nigrum* | 268 / 273 | 28.4 / 38.3 | +9.9 | 6.2 | 0.001 | 0.017 |
| *Juniperus communis* | 22 / 39 | 0.1 / 0.6 | +0.5 | 2.3 | 0.007 | 1 |
| *Pedicularis lapponica* | 56 / 39 | 0.1 / + | -0.1 | -2.2 | 0.005 | 1 |
| *Pinus sylvestris* | 2 / 11 | + / + | + | 1.9 | 0.001 | 0.173 |
| *Vaccinium myrtillus* | 147 / 175 | 13.4 / 9.1 | -4.3 | -4.1 | 0.001 | 0.017 |
| *Vaccinium uliginosum* | 171 / 177 | 2.3 / 4.5 | +2.2 | 4.5 | 0.001 | 0.032 |
| *Vaccinium vitis-idaea* | 253 / 253 | 3.3 / 2.4 | -0.9 | -2.7 | 0.003 | 0.428 |
| Bryophytes |  |  |  |  |  |  |
| *Aulacomnium palustre* | 17 / 9 | 0.3 / + | -0.3 | -1.5 | 0.003 | 0.385 |
| *Hepaticae* | 184 / 160 | 2.4 / 0.9 | -1.5 | -3.3 | 0.001 | 0.017 |
| *Pleurozium schreberi* | 162 / 189 | 7.8 / 12.5 | +4.7 | 3.5 | 0.001 | 0.064 |
| *Polytrichum spp.* | 206 / 197 | 2.3 / 1.1 | -1.2 | -3.2 | 0.001 | 0.032 |
| Lichens |  |  |  |  |  |  |
| *Cetraria spp.* | 201 / 178 | 0.8 / 0.4 | -0.3 | -3.0 | 0.001 | 0.079 |
| *Cladonia arbuscula* | 232 / 233 | 2.7 / 1.6 | -1.1 | -3.0 | 0.002 | 0.310 |
| *Cladonia rangiferina* | 144 / 205 | 0.3 / 0.7 | +0.4 | 3.6 | 0.001 | 0.017 |
| *Cladonia stellaris* | 72 / 83 | 0.6 / 0.2 | -0.4 | -2.3 | 0.004 | 0.532 |
| *crustaceous lichens* | 169 / 136 | 2.3 / 1.1 | -1.1 | -2.3 | 0.001 | 0.203 |
| *Stereocaulon spp.* | 134 / 125 | 6.2 / 1.0 | -5.2 | -5.6 | 0.001 | 0.017 |

## Discussion

Fennoscandian oligotrophic heathland and tundra plant communities have become compositionally more similar during the past five decades when compared to their reference condition in 1964–75. The observed homogenization pattern is not dependent on the biogeographical zonation or the degree of continentality-humidity (excluding subcontinental conditions), suggesting a distinct and pervasive trend in the study area. Our results indicate that the observed large-scale homogenization is resulting from a compositional converge of plant communities among different habitat types and is linked to the increased abundance of evergreen dwarf shrub *E. nigrum.* Lichen communities showed increased similarity at many sites, whereas bryophyte communities seemed to be more stable, revealing differing patterns of long-term diversity dynamics among these growth forms. Our findings extend the observational evidence of biotic homogenization pattern found in Bråthen et al. (2024) to a wider area and species pool in northern European heathlands and tundra and align with the predictions of tundra vegetation homogenization resulting from climate change (Steward et al. 2018, Niittynen et al. 2020). The Fennoscandian tundra is transitioning from a landscape of distinct habitat types towards one dominated by *E. nigrum*. The observed ecosystem-wide homogenization likely poses a threat to ecosystem multifunctionality (van der Plas et al. 2016, Hautier et al. 2017).

### The observed patterns of compositional change

The observed homogenization across the whole study area occurred within all growth forms, most clearly within vascular plant and lichen communities. The increased similarity within vascular plant communities was strong across all biogeographical zones and continentality-humidity classes (except subcontinental), but responses of other growth forms were more varied. The increased similarity of bryophyte communities was limited to the northern boreal zone, where the common boreal species, *Pleurozium schreberi*, doubled its cover over time (Table S7). The studied northern boreal sites locate at or near the treeline and such expansion of boreal species into treeless heaths may indicate ongoing borealization (see also, Vuorinen et al. 2017; Villén-Peréz et al. 2020). The observed homogenization of lichen communities in the hemioroarctic and oroarctic zones coincided with a drastic temporal decline in the cover of *Stereocaulon* spp. and crustaceous lichens (Table S7), which prefer open ground. While herbivory has clearly been shown to have a negative effect on lichen cover and biomass (Kumpula et al. 2014, Olofsson et al. 2014, Bernes et al. 2015), such strong species-specific changes suggest that the increased abundance of vascular plants, or shrub encroachment, may also drive lichen decline (Cornelissen et al. 2001). Subcontinental conditions, in turn, may be related to the stability in compositional dissimilarity over time, but this may simply be due to the smaller sample size and fewer species showing significant change (Table S8).

The compositional characteristics of previously more distinct habitat types have become blurred over time (see also, Ross et al. 2012), leading to a landscape-scale homogenization. The number of plots in different habitat types reflect the real share of types in the study area (Haapasaari, 1988), strengthening the generalizability of the observed landscape-scale homogenization across the Fennoscandian heathlands and tundra. Pronounced compositional shifts had occurred especially in *V. myrtillus-*, *C. vulgaris-* and in windswept types (Fig. 2d), all of which had experienced a notable *E. nigrum* encroachment over time (Fig. 4). In *V. myrtillus-*type, *E. nigrum* had replaced *V. myrtillus* as a dominant species, which was also reflected in the increased evenness of vascular plants. The competitive ability of *B. nana* likely prevented the expansion of *E. nigrum* into the *B. nana-*type. Interestingly, the evenness of vascular plants had also increased in *E. nigrum-*type, where deciduous shrubs had increased in cover (Fig. 4). Simultaneously, the mean cover of *E. nigrum* had remained stable and relatively high (c. 45%) over time, but especially *V. myrtillus* had clearly established in new plots (Table S9). Even though the seedlings of *V. myrtillu*s are negatively affected by the allelopathy (Pilsbacher et al. 2021), seeds can germinate and establish through reindeer faeces (Bråthen et al. 2007).

The more even dominance of dwarf shrubs in the resurvey likely contributed to the unchanged dissimilarity within vascular plant communities in many habitat types and the heterogenization of vascular plant communities within *V. myrtillus-*type. In this habitat type, the cover of the originally most dominant species, *V. myrtillus*, became partially replaced by other species. It should be closely monitored whether this development and changes in dominance patterns are transitory or more lasting. In contrast, we observed homogenization of lichen communities in three common habitat types (*E. nigrum-, V. myrtillus-* and *B. nana-*types) while lichen evenness increased only in *E. nigrum-*type. Despite a general decrease in lichen cover, their evenness increased also in certain conditions (hemiarctic to oroarctic and subcontinental to indifferent), meaning that the current taxa are now more uniformly abundant, albeit covering less ground overall. To conclude, our results suggest that the process through which the homogenization occurs in the study area is not through species losses (decreasing species richness) but through abundance change of dominant species, especially of vascular plants (e.g. *E. nigrum*) and lichens (e.g. *Stereocaulon* spp.).

### Potential causes and consequences of the observed compositional changes

We did not test the effect of environmental drivers on the observed homogenization patterns, but several lines of evidence suggest that changes in snow cover—in particular, decreased snow cover duration— may play a key role in the compositional changes (Bokhorst et al. 2016, Niittynen et al. 2018), comparable to observations in many snow-dependent habitat types (Morgan & Walker 2023, Choler et al. 2024). *E. nigrum* generally performs better under changing snow cover conditions (Bienau et al. 2014) and is relatively tolerant to seasonal and inter-annual climatic variability (González et al. 2019) compared to many other species, such as *V. myrtillus* that needs more snow to thrive. Earlier snowmelt and spring exposure have been shown to advance the onset of shoot growth in *E. nigrum*, while delaying shoot growth in *V. myrtillus* (Wipf et al. 2009). This may have facilitated *E. nigrum* to gain more ground over time, especially in habitats originally dominated by *V. myrtillus*. The direct response of evergreen dwarf shrubs to summer warming is generally not as strong as the response of low-growing and tall deciduous shrubs (Elmendorf et al. 2012). Warmer temperatures have most likely enhanced the growth of low-growing deciduous shrubs in the study area (e.g. *B. nana,* Table 2), similar to other areas across the circumpolar Arctic (Elmendorf et al. 2012). However, the larger increases are controlled by reindeer grazing (Olofsson et al. 2009, Bråthen et al. 2017, Ramirez et al. 2024) that, in turn, do not profoundly inhibit or promote *E. nigrum* expansion (Vowles et al. 2017, Tuomi et al. 2024). However, the effects of long-term snow cover change on the observed compositional changes need further studies.

In the time of the original survey, habitat types with more spatial and compositional variation, each dominated by different species, likely supported the provision of multiple differing functions across spatial scales (Isbell et al. 2011, Mori et al. 2018). Hence, the observed homogenization and eased local dominance have likely reduced ecosystem multifunctionality of the Fennoscandian tundra on a regional scale (Hautier et al. 2017, Liu et al. 2021). Fennoscandian tundra is moving away from the state where the functions and processes were shaped by distinct habitat types and several dominant species within landscapes, towards the ecosystem functioning shaped by *E. nigrum-*dominated communities. The observed taxonomic homogenization of Fennoscandian tundra may have led to functional homogenization, whereby communities have become similar not only by their species composition but also by their functional trait combination and variation (Hillebrand et al. 2008). Traits of *E. nigrum* have become overrepresented in communities across the ecosystem, hence the related ecosystem functions they provide have increased. For instance, *E. nigrum* dominance harms the growth (Bråthen et al. 2010, González et al. 2015) and reproduction (Pilsbacher et al. 2021) of boreal and tundra species and inhibits nutrient cycles (Tybirk et al. 2000). *E. nigrum* expansion has also been shown to affect functions such as reindeer pasture quality (Tuomi et al. 2024), which is further impacted by the observed decrease in lichen cover and diversity. In general, the effects of *E. nigrum* expansion on multifunctionality should be addressed swiftly, as recent studies indicate that its encroachment has the potential to accelerate (Maliniemi et al. 2024, Tuomi et al. 2024). Understanding the influence of different shrub species on long-term compositional changes in the boreal-Arctic ecosystems is crucial, and we argue along with Vowles & Björk (2019) that if the impacts of shrub expansion to tundra vegetation is being evaluated based only on the characteristics of deciduous shrub species, our understanding of the ecosystem feedback resulting from vegetation shifts will be inadequate.

Our study highlights the importance of considering compositional and beta diversity changes when assessing the condition of habitat types and ecosystems (e.g., Kontula & Raunio 2019, Pedersen et al. 2021), as the measures of alpha diversity (especially species richness) alone dot not necessarily capture all the fundamental shifts in the communities (see also, Mori et al. 2018). Essentially, both local and regional responses over time must be considered (Li et al. 2020, Dornelas et al. 2023) as biodiversity effects are often experienced at different spatial scales, as emphasized by our findings of regional-scale homogenization compared to no change or even heterogenization at the level of habitat types. Lastly, our findings highlight the importance of large-scale and long-term monitoring of common species and habitat types not yet endangered, which may prove to be blind spots that threaten biodiversity, as argued by Tuomi et al. (2024).

## Author contributions

TM initiated the study. TM wrote the first draft with a major contribution from PK. TM, JK, TBR and RV collected the data. TM and PK analyzed the data. All authors edited and commented on the manuscript throughout the process.

## Acknowledgements

We thank Anu Skog, Emmi Virsula and Elina Nystedt for assisting in the resurvey. TM was funded by Foundations Post Doc Pool (Finnish Cultural Foundation) and Biodiverse Anthropocenes research project supported by University of Oulu & The Research Council of Finland Profi6 336449. PK acknowledges funding from Kvantum Institute of University of Oulu, KAB from the Norwegian Research Council project # 302749, RV from Academy of Finland project # 259072 and JK from INTERACT (International Network for Terrestrial Research and Monitoring in the Arctic) grant numbers 262693 and 730938.

## Notes

### Competing Interest Statement

The authors have declared no competing interest.

## References

Alfonsi, E., Benot M.-L., Fievet, V. and Alard, D. 2017 Addressing species turnover and community changes in vegetation resurvey studies. Appl. Veg. Sci. 20: 172–182.

Anderson, M. J., Ellingsen, K. E. and McArdle, B. H. 2006. Multivariate dispersion as a measure of beta diversity. Ecol. Lett. 9: 683–693.

Asplund, J. and Wardle, D. A. 2017. How lichens impact on terrestrial community and ecosystem properties. Biol. Rev. 92: 1720–1738.

Bernes, C., Bråthen, K. A., Forbes, B. C., Speed, J. D. M. and Moen, J. 2015. What are the impacts of reindeer/caribou (Rangifer tarandus L.) on arctic and alpine vegetation? A systematic review. Environ. Evid. 4: 4.

Bienau M. J., Hattermann, D., Kröncke, M., Kretz, L., Otte, A., Eiserhardt, W. L., Milbau, A., Graae, B. J., Durka, W. and Eckstein, R. L. 2014. Snow cover consistently affects growth and reproduction of *Empetrum hermaphroditum* across latitudinal and local climatic gradients. Alpine Bot. 124: 115–129.

Bokhorst, S., Pedersen, S. H., Brucker, L., Anisimov, O., Bjerke, J. W., Brown, R. D., Ehrich, D., Essery, R. L. H., Heilig, A., Ingvander, S., Johansson, C., Johansson, M., Jónsdóttir, I. S., Inga, N., Luojus, K., Macelloni, G., Mariash, H., McLennan, D., Rosqvist, G. N., Sato, A., Savela, H., Schneebeli, M., Sokolov, A., Sokratov, S. A., Terzago, S., Vikhamar-Schuler, D., Williamson, S., Qiu, Y. and Callaghan, T. V. 2016. Changing Arctic snow cover: A review of recent developments and assessment of future needs for observations, modelling, and impacts. Ambio 45: 516–537.

Bråthen, K. A., González, V. T., Iversen, M., Killengreen, S., Ravolainen, V. T., Ims, R. A. and Yoccoz, N. G. 2007. Endozoochory varies with ecological scale and context. Ecography 30: 308–320.

Bråthen K. A., Fodstad C. H., and Gallet C. 2010. Ecosystem disturbance reduces the allelopathic effects of Empetrum hermaphroditum humus on tundra plants. J. Veg. Sci. 21: 786–795.

Bråthen, K. A. and Ravolainen, V. T. 2015. Niche construction by growth forms is as strong a predictor of species diversity as environmental gradients. J. Ecol. 103: 701–713.

Bråthen, K. A., Ravolainen, V. T., Stein, A., Tveraa, T. and Ims, R. A. 2017. Rangifer management controls a climate-sensitive tundra state transition. Ecol. Appl. 27: 2416–2427.

Bråthen, K. A., Gonzalez, V. T. and Yoccoz, N. G. 2018. Gatekeepers to the effects of climate warming? Niche construction restricts plant community changes along a temperature gradient. Perspect. Plant Ecol. 30: 71–81.

Bråthen, K. A., Tuomi, M., Kapfer, J., Böhner, H. and Maliniemi, T. 2024. Changing species dominance patterns of Boreal-Arctic heathlands: evidence of biotic homogenization. Ecography 2024: e07116.

CBD (2022). Convention on Biological Diversity. Kunming-Montreal Global Biodiversity Framework. - <www.cbd.int.>.

Choler, P., Bayle, A., Fort, N. and Gascoin, S. 2024. Waning snowfields have transformed into hotspots of greening within the alpine zone. Nat. Clim. Change. In press. 10.1038/s41558-024-02177-x

Cornelissen, J. H. C, Callaghan, T. V., Alatalo, J. M., Michelsen, A., Graglia, E., Hartley, A. E., Hik, D. S., Hobbie, S. E., Press, M. C., Robinson, C. H., Henry, G. H. R., Shaver, G. R., Phoenix, G. K., Gwynn Jones, D., Jonasson, S., Chapin III, F. S., Molau, U., Neill, C., Lee, J. A., Melillo, J. M., Sveinbjörnsson, B. and Aerts, R. 2001. Global change and arctic ecosystems: is lichen decline a function of increases in vascular plant biomass? J. Ecol. 89: 984–994.

Cornelissen, J. H. C., Lang, S. I., Soudzilovskaia, N. A. and During, H. J. 2007. Comparative Cryptogam Ecology: A Review of Bryophyte and Lichen Traits that Drive Biogeochemistry. Ann. Bot. 99: 987–1001.

Dornelas, M., Chase, J. M., Gotelli, N. J., Magurran, A. E., McGill, B. J., Antão, L. H., Blowes, S. A., Daskalova, G. N., Leung, B., Martins, I. S., Moyes, F., Myers-Smith, I. H., Thomas, C. D., and Vellend, M. 2023. Looking back on biodiversity change: lessons for the road ahead. Philos. Trans. R. Soc. B 378: 20220199.

Dray, S., Bauman, D., Blanchet, G., Borcard, D., Clappe, S., Guenard, G., Jombart, T., Larocque, G., Legendre, P., Madi, N., Wagner, H. H. and Siberchicot, A. 2024. adespatial: Multivariate Multiscale Spatial Analysis – <https://CRAN.R-project.org/package=adespatial>.

Elmendorf, S. C., Henry, G. H. R., Hollister, R. D., Björk, R. G., Boulanger-Lapointe, N., Cooper, E. J., Cornelissen, J. H. C., Day, T. A., Dorrepaal, E., Elumeeva, T. G., Gill, M., Gould, W. A., Harte, J., Hik, D. S., Hofgaard, A., Johnson, D. R., Johnstone, J. F., Jónsdóttir, I. S., Jorgenson, J. C., Klanderud, K., Klein, J. A., Koh, S., Kudo, G., Lara, M., Lévesque, E., Magnússon, B., May, J. L., Mercado-Dı‘az, J. A., Michelsen, A., Molau, U., Myers-Smith, I. H., Oberbauer, S. F., Onipchenko, V. G., Rixen, C., Schmidt, N. M., Shaver, G. R., Spasojevic, M. J., Þórhallsdóttir, Þ. E., Tolvanen, A., Troxler, T., Tweedie, C. E., Villareal, S., Wahren, C.-H., Walker, X., Webber, P. J., Welker, J. M. and Wipf, S. 2012. Plot-scale evidence of tundra vegetation change and links to recent summer warming. *Nat*. Clim. Change 2: 453–457.

Finderup Nielsen, T., Sand-Jensen, K., Dornelas, M. and Bruun, H. H. 2019. More is less: net gain in species richness, but biotic homogenization over 140 years. Ecol. Lett. 22: 1650–1657.

García Criado, M., Myers-Smith, I. H, Bjorkman, A. D., Elmendorf, S. C., Normand, S., Aastrup, P., Aerts, R., Alatalo, J. M., Baeten, L., Björk, R. G., Björkman, M. P., Boulanger-Lapointe, N., Butler, E. E., Cooper, E. J., Cornelissen, J. H. C., Daskalova, G. N., Henry, G. H. R., Hollister, R. D., Høye, T. T., Fadrique, B., Jacobsen, I. B. D., Jägerbrand, A. K, Jónsdóttir, I. S., Kaarlejärvi, E., Khitun, O., Klanderud, K., Kolari, T. H. M., Lang, S. I., Lecomte, N., Lenoir, J., Macek, P., Messier, J., Michelsen, A., Molau, U., Muscarella, R., Nielsen, M.-L., Petit Bon, M., Post, E., Raundrup, K., Rinnan, R., Rixen, C., Ryde, I., Serra-Diaz, J. M., Schaepman-Strub, G., Schmidt, N. M., Schrodt, F., Sjögersten, S., Steinbauer, M. J., Stewart, L., Strandberg, B., Tolvanen, A., Tweedie, C. E. and Vellend, M. 2024. Plant diversity dynamics over space and time in a warming Arctic. Preprint in EcoEvoRxiv: 10.32942/X2MS4N

González, V. T., Junttila, O., Lindgård, B., Reiersen, R., Trost, K. and Bråthen, K. A. 2015. Batatasin-III and the allelopathic capacity of Empetrum nigrum. Nord. J. Bot. 33: 225–231.

González, V. T., Moriana-Armendariz, M., Hagen, S. B., Lindgård, B., Reiersen, R. and Bråthen, K. A. 2019. High resistance to climatic variability in a dominant tundra shrub species. PeerJ 7: e6967.

Haapasaari, M. 1988. The oligotrophic heath vegetation of northern Fennoscandia and its zonation. Acta Bot. Fenn. 135: 1–129.

Hautier, Y., Isbell, F., Borer, E. T., Seabloom, E. W., Harpole, W. S., Lind, E. M., MacDougall, A. S., Stevens, C. J., Adler, P. B., Alberti, J., Bakker, J. D., Brudvig, L. A., Buckley, Y. M., Cadotte, M., Caldeira, M. C., Chaneton, E. J., Chu, C., Daleo, P., Dickman, C. R., Dwyer, J. M., Eskelinen, A., Fay, P. A., Firn, J., Hagenah, N., Hillebrand, H., Iribarne, O., Kirkman, K. P., Knops, J. M. H., La Pierre, K. J., McCulley, R. L., Morgan, J. W., Pärtel, M., Pascual, J., Price, J. N., Prober, S. M., Risch, A. C., Sankaran, M., Schuetz, M., Standish, R. J., Virtanen, R., Wardle, G. M., Yahdjian, L. and Hector, A. 2018. Local loss and spatial homogenization of plant diversity reduce ecosystem multifunctionality. *Nat*. Ecol. Evol 2: 50–56.

Herve, M. 2023. RVAideMemoire: Testing and Plotting Procedures for Biostatistics. –<https://CRAN.R-project.org/package=RVAideMemoire>.

Hillebrand, H., Bennett, D. M. and Cadotte, M. W. 2008. Consequences of Dominance: a Review of Evenness Effects on Local and Regional Ecosystem Processes. Ecology 89: 1510–1520.

IPBES. 2019. Global assessment report on biodiversity and ecosystem services of the Intergovernmental Science-Policy Platform on Biodiversity and Ecosystem Services. Brondizio, E. S., Settele, J., Díaz, S. and H. T. Ngo (eds). IPBES secretariat, Bonn, Germany. 1148 pages.

Isbell, F., Calcagno, V., Hector, A., Connolly, J., Harpole, W. S., Reich, P. B., Scherer-Lorenzen, M., Schmid, B., Tilman, D., van Ruijven, J., Weigelt, A., Wilsey, B. J., Zavaleta, E. S. and Loreau, M. 2011. High plant diversity is needed to maintain ecosystem services. Nature 477: 199–202.

Kapfer, J., Hédl, R., Jurasinski, G., Kopecký, M., Schei, F. H. and Grytnes, J.-A. 2017. Resurveying historical vegetation data – opportunities and challenges. Appl. Veg. Sci. 20: 164–171.

Kontula, T. and Raunio, A. 2019. Threatened Habitat types in Finland 2018. Red List of Habitats – results and basis for Assessment. The Finnish Environment 2/2019. Finnish Environment Institute and Ministry of the Environment, Helsinki. 254 pages.

Kumpula, J., Kurkilahti, M., Helle, T. and Colpaert, A. 2014. Both reindeer management and several other land use factors explain the reduction in ground lichens (Cladonia spp.) in pastures grazed by semi-domesticated reindeer in Finland. Reg. Environ. Change 14: 541–559.

Kuusisto, I., Huttunen, S. and Virtanen, R. 2024. Tundra plant communities along the mesotopographic gradient in NE Finland. Nord. J. Bot. In press. 10.1111/njb.04430

Legendre, P. 2019. A temporal beta-diversity index to identify sites that have changed in exceptional ways in space-time surveys. Ecol. Evol. 9: 3500–3514.

Lett, S., Jónsdóttir, I. S., Becker-Scarpitta, A., Christiansen, C. T., During, H., Ekelund, F., Henry, G. H. R., Lang, S., Michelsen, A., Rousk, K., Alatalo, J., Betway, K. R., Busca, S., Callaghan, T., Carbognani, M., Cooper, E. J., Cornelissen, J. H. C., Dorrepaal, E., Egelkraut, D., Elumeeva, T. G., Haugum, S. V., Hollister, R. D., Jägerbrand, A. K., Keuper, F., Klanderud, K., Lévesque, E., Liu, X., May, J., Michel, P., Mörsdorf, M., Petraglia, A., Rixen, C., Robroek, B. J. M., Rzepczynska, A. Soudzilovskaia, N. A., Tolvanen, A., Vandvik, V., Volkov, I., Volkova, I. and van Zuijlen, K. 2021. Can bryophyte groups increase functional resolution in tundra ecosystems? Arct. Sci. 8: 609–637.

Li, D., Olden J. D., Lockwood J. L., Record S., McKinney, M. L. and Baiser, B. 2020. Changes in taxonomic and phylogenetic diversity in the Anthropocene. Proc. R. Soc. B. 287: 20200777

Liu, D., Chang, P.-H. S., Power, S. A., Bell, J. N. B. and Manning, P. 2021. Changes in plant species abundance alter the multifunctionality and functional space of heathland ecosystems. New Phytol. 232: 1238–1249.

Maliniemi, T., Kapfer, J., Saccone, P., Skog, A. and Virtanen, R. 2018. Long-term vegetation changes of treeless heath communities in northern Fennoscandia: links to climate change trends and reindeer grazing. J. Veg. Sci. 29: 469–479.

Maliniemi, T., Happonen, K. and Virtanen. R. 2019. Site fertility drives temporal turnover of vegetation at high latitudes. Ecol. Evol. 9: 13255–13266.

Maliniemi, T., Lohi, J., Alahuhta, J., Huusko, K. and Virtanen, R. 2024. Dwarf shrub expansion and loss of lichens distinctly dominate multi-decadal changes in northern boreal understory plant communities. Nord. Geogr. Publ. Accepted manuscript. 10.30671/nordia.145998

McKinney, M. L. and Lockwood, J. L. 1999. Biotic homogenization: a few winners replacing many losers in the next mass extinction. Trends Ecol. Evol. 14: 450–453.

McCune, J. L. and Vellend, M. 2013. Gains in native species promote biotic homogenization over four decades in a human-dominated landscape. J. Ecol. 101: 1542–1551.

Mod, H. K., Heikkinen, R. K., le Roux, P. C., Wisz, M. S. and Luoto, M. 2016. Impact of biotic interactions on biodiversity varies across a landscape. J. Biogeogr. 43: 2412–2423.

Morgan, J. and Walker, Z. 2023. Early-melting snowpatch plant communities are transitioning into novel states. Sci. Rep. 13: 16520.

Mori, A. S., Isbell, F. and Seidl, R. 2018. β-Diversity, Community Assembly, and Ecosystem Functioning. Trends Ecol. Evol. 33: 549–564.

Myers-Smith, I. H., Kerby, J. T., Phoenix, G. K., Bjerke, J. W., Epstein, H. E., Assmann, J. J. Christian John, Andreu-Hayles, L., Angers-Blondin, S., Beck, P. S. A., Berner, L. T., Bhatt, U. S., Bjorkman, A. D., Blok, D., Bryn, A., Christiansen, C. T., Cornelissen, J. H. C., Cunliffe, A. M., Elmendorf, S. C., Forbes, B. C. Goetz, S. J., Hollister, R. D., de Jong, R., Loranty, M. M., Macias-Fauria, M., Maseyk, K., Normand, S., Olofsson, J., Parker, T. C., Parmentier, F.-J. W., Post, E., Schaepman-Strub, G., Stordal, F., Sullivan, P. F., Thomas, H. J. D., Tømmervik, H., Treharne, R., Tweedie, C. E., Walker, D. A., Wilmking, M. and Wipf, S. 2020. Complexity revealed in the greening of the Arctic. *Nat*. Clim. Change 10: 106–117.

Niittynen, P., Heikkinen, R. K. and Luoto, M. 2018. Snow cover is a neglected driver of Arctic biodiversity loss. *Nat*. Clim. Change 8: 997–1001.

Niittynen, P., Heikkinen, R. K. and Luoto, M. 2020. Decreasing snow cover alters functional composition and diversity in the tundra. Proc. Natl. Acad. Sci. U.S.A. 117: 21480–21487.

Oksanen, J., Simpson, G. L., Blanchet, F. G., Kindt, R., Legendre, P., Minchin, P. R., O’Hara, R. B., Solymos, P., Stevens, M. H. H, Szoecs, E., Wagner, H., Barbour, M., Bedward, M., Bolker, B., Borcard, D., Carvalho, G., Chirico, M., De Caceres, M., Durand, S., [aut], Evangelista, H. B. A., FitzJohn, R., Friendly, M., Furneaux, B., Hannigan, G., Hill, M. O., Lahti, L., McGlinn, D., Ouellette, M.-H., Ribeiro Cunha, E., Smith, T., Stier, A., Ter Braak, C. J. F., Weedon, J. 2024. vegan: Community Ecology Package. – <https://CRAN.R-project.org/package=vegan>.

Olden, J. D., Poff, N. L., Douglas, M. R., Douglas, M. E. and Fausch, K. D. 2004. Ecological and evolutionary consequences of biotic homogenization. Trends Ecol. Evol. 19: 18–24.

Olden, J. D. and Rooney T. P. 2006. On defining and quantifying biotic homogenization. Global Ecol. Biogeog. 15: 113–120.

Olofsson, J., Oksanen, L., Callaghan, T., Hulme, P. E., Oksanen T. and Suominen, O. 2009. Herbivores inhibit climate-driven shrub expansion on the tundra. Global Change Biol. 15: 2691–2693.

Olofsson, J., Oksanen, L., Oksanen, T., Tuomi, M., Hoset, K. S., Virtanen, R. and Kyrö, K. 2014. Long-term experiments reveal strong interactions between lemmings and plants in the Fennoscandian highland tundra. Ecosystems 17: 606–615.

Pedersen, Å. Ø., Jepsen, J. U., Paulsen, I. M. G., Fuglei, E., Mosbacher, J. B., Ravolainen, V., Yoccoz, N. G., Øseth, E., Böhner, H., Bråthen, K. A., Ehrich, D., Henden, J.-A., Isaksen, K., Jakobsson, S., Madsen, J., Soininen, E., Stien, A., Tombre, I., Tveraa, T., Tveito, O. E., Vindstad, O. P. L. and Ims, R. A. 2021. Norwegian Arctic Tundra: a Panel-based Assessment of Ecosystem Condition. In: Report series, vol. 153. Norwegian Polar Institute.

Pellissier, L., Bråthen, K. A., Pottier, J., Randin, C. F., Vittoz, P., Dubuis, A., Yoccoz, N. G., Alm, T., Zimmermann, N. E. and Guisan, A. 2010. Species distribution models reveal apparent competitive and facilitative effects of a dominant species on the distribution of tundra plants. Ecography 33: 1004–1014.

Pilsbacher, A. K., Lindgård, B., Reiersen, R., Gonzalez, V. and Bråthen, K. A. 2021. Interfering with neighbouring communities: Allelopathy astray in the tundra delays seedling development. Funct. Ecol. 35: 266–276.

Post, E. (2013). Erosion of community diversity and stability by herbivore removal under warming. Proc. R. Soc. B 280: 20122722.

Ramirez, J. I., Sundqvist, M., Lindén, E., Björk, R. G., Forbes, B. C., Suominen, O., Tyler, T., Virtanen, R. and Olofsson, J. 2024. Reindeer grazing reduces climate-driven vegetation changes and shifts trophic interactions in the Fennoscandian tundra. Oikos 2024: e10595.

Rantanen, M., Karpechko, A. Y., Lipponen, A., Nordling, K., Hyvärinen, O., Ruosteenoja, K., Vihma, T. and Laaksonen, A. 2022. The Arctic has warmed nearly four times faster than the globe since 1979. Commun. Earth Environ. 3: 168.

Rantanen, M., Kämäräinen, M., Niittynen, P., Phoenix, G. K., Lenoir, J., Maclean, I., Luoto, M. and Aalto, J. 2023. Bioclimatic atlas of the terrestrial Arctic. Sci. Data 10: 40.

Ravolainen, V. T., Yoccoz, N. G., Bråthen, K. A., Ims, R. A., Iversen, M. and González, V. T. 2010. Additive partitioning of diversity reveals no scale-dependent impacts of large ungulates on the structure of tundra plant communities. Ecosystems 13: 157–170.

Rooney, T. 2009. High white-tailed deer densities benefit graminoids and contribute to biotic homogenization of forest ground-layer vegetation. Plant Ecol. 202: 103–111.

Ross, L. C., Woodin, S. J., Hester, A. J., Thompson, D. B. A. and Birks, H. J. B. 2012. Biotic homogenization of upland vegetation: patterns and drivers at multiple spatial scales over five decades. J. Veg. Sci. 23: 755–770.

Socolar, J. B., Gilroy, J. J., Kunin, W. E. and Edwards, D. P. 2016. How Should Beta-Diversity Inform Biodiversity Conservation? Trends Ecol. Evol. 31: 67–80.

Soininen, E., Magnusson, M., Jepsen, J., Eide, N., Yoccoz, N. G., Angerbjörn, A., Breisjøberget, J. I., Ecke, F., Ehrich, D., Framstad, E., Henttonen, H., Hörnfeldt, B., Killengren, S., Olofsson, J., Oksanen, L., Oksanen, T., Tveito, O and Ims, R. 2024. Macroecological patterns of rodent population dynamics shaped by bioclimatic gradients. Ecography. Accepted manuscript. Preprint: https://www.techrxiv.org/doi/full/10.22541/au.169211023.35851530

Stark, S, Ylänne, H. and Kumpula, J. 2021. Recent changes in mountain birch forest structure and understory vegetation depend on the seasonal timing of reindeer grazing. J. Appl. Ecol. 58: 941–952.

Stark, S., Horstkotte, T., Kumpula, J., Olofsson, J., Tømmervik, H. and Turunen, M. 2023. The ecosystem effects of reindeer (Rangifer tarandus) in northern Fennoscandia: Past, present and future. *Perspect*. Plant Ecol. 58: 125716.

Stewart, L., Simonsen, C. E., Svenning, J.-C., Schmidt, N. M. and Pellissier, L. 2018. Forecasted homogenization of high Arctic vegetation communities under climate change. J. Biogeogr. 45: 2576–2587.

Turetsky, M. R., Bond-Lamberty, B., Euskirchen, E., Talbot, J., Frolking, S., McGuire, A. D. and Tuittila, E. S. 2012. The resilience and functional role of moss in boreal and arctic ecosystems. New Phytol. 196: 49–67.

Tuomi, M. W., Utsi, T. A., Yoccoz, N. G., Armstrong, C. V., Gonzales, W., Hagen, S. B., Jónsdóttir I. S., Pugnaire, F. I., Shea, K., Wardle, D., A. Zielosko and Bråthen, K. A. 2024. The increase of an allelopathic and unpalatable plant undermines reindeer pasture quality and current management in the Norwegian tundra. Commun. Earth Environ. 5: 414.

Tybirk, K., Nilsson, M.-C., Michelsen, A., Kristensen, H. L., Shevtsova, A., Strandberg, M. T., Johansson, M., Nielsen, K. E., Riis-Nielsen, T., Stranberg, B. and Johnsen, I. 2000. Nordic *Empetrum* dominated ecosystems: Function and susceptibility to environmental changes. Ambio 29: 90–97.

Vellend, M., Baeten, L., Becker-Scarpitta, A., Boucher-Lalonde, V., McCune, J. L., Messier, J., Myers-Smith, I. H. and Sax, D. F. 2017. Plant biodiversity change across scales during the Anthropocene. Ann. Rev. Plant Biol. 68: 563–586.

Villén-Peréz, S., Heikkinen, J., Salemaa, M. and Mäkipää, R. 2020. Global warming will affect the maximum potential abundance of boreal plant species. Ecography 43: 801–811.

Virtanen, R., Oksanen, L., Oksanen, T., Cohen, J., Forbes, B. C., Johansen, B., Käyhkö, J., Olofsson, J., Pulliainen, J. and Tømmervik, H. 2016. Where do the treeless tundra areas of northern highlands fit in the global biome system: Towards an ecologically natural subdivision of the tundra biome. Ecol. Evol. 6: 143–158.

Vowles, T., Gunnarsson, B., Molau, U., Hickler, T., Klemedtsson, L. and Björk, R. G. 2017. Expansion of deciduous tall shrubs but not evergreen dwarf shrubs inhibited by reindeer in Scandes mountain range. J. Ecol. 105: 1547–1561.

Vowles, T. and Björk, R. G. 2019. Implications of evergreen shrub expansion in the Arctic. J. Ecol. 107: 650–655.

Vuorinen, K. E. M., Oksanen, L., Oksanen, T., Pyykönen, A., Olofsson, J. and Virtanen, R. 2017. Open tundra persist, but arctic features decline—Vegetation changes in the warming Fennoscandian tundra. Global Change Biol. 23: 3794–3807.

Walker, M. D., Wahren, C. H., Hollister, R. D., Henry, G. H. R. R., Ahlquist, L. E., Alatalo, J. M., Bret-Harte, M. S., Calef, M. P., Callaghan, T. v., Carroll, A. B., Epstein, H. E., Jónsdóttir, I. S., Klein, J. A., Magnússon, B., Molau, U., Oberbauer, S. F., Rewa, S. P., Robinson, C. H., Shaver, G. R., Suding, K. N., Thompson, C. C., Tolvanen, A., Totland, Ø., Turner, P. L., Tweedie, C. E., Webber, P. J. and Wookey, P. A. 2006. Plant community responses to experimental warming across the tundra biome. Proc. Natl. Acad. Sci. U.S.A. 103: 1342–1346.

Wilson, S. D. and Nilsson, C. 2009. Arctic alpine vegetation change over 20 years. Global Change Biol. 15: 1676–1684.

Wipf, S., Stoeckli, V. and Bebi, P. 2009. Winter climate change in alpine tundra: plant responses to changes in snow depth and snowmelt timing. Climatic Change 94: 105–121.

