## Supplementary Material for "Long-term homogenization of vascular plant and lichen communities across Fennoscandian heathlands and tundra is connected to the expansion of an allelopathic dwarf shrub"

**Table S1.** Species that were merged.

| **Vascular plants** | **Include** |
|---|---|
| *Agrostis ssp.* | *A. capillaris, A. mertensii, A. sp* |
| *Poa ssp.* | *P. alpigena, P. sp* |

| **Bryophytes** | **Include** |
|---|---|
| *Aulacomnium spp.* | *A. palustere, A. turgidum* |
| *Dicranella spp.* | *D. sp* |
| *Dicranum spp.* | *D. drummondii, D. elongatum, D. fuscescens, D. flexicaule, D.majus, D. montanum, D. polysetum, D. scoparium, D. spurium, D. undulatum, D. sp* |
| *Ditrichum spp.* | *D. flexicaule, D. zonatum* |
| *Kiaeria spp.* | *K. blyttii, K. gracilis* |
| *Mnium spp.* | *M. hornum, M. spinosum, M. thomsonii* |
| *Plagiothecium spp.* | *P. denticulatum, P. laetum* |
| *Pogonatum spp.* | *P. dentatum, P.urnigerum* |
| *Pohlia spp.* | *P. nutans, P. cruda. P.sp* (most often *P. nutans*) |
| *Polytrichum spp.* | *P. commune, P. hyperboreum, P. juniperinum, P. piliferum, P. strictum, Polytrichastrum alpinum, P. sexangulare* |
| *Ptychostomum spp.* | *P. sp* |
| *Racomitrium spp.* | *R. lanoginosum, R. microcarpon, R. fasciculare* |
| *Rhytidiadelphus spp.* | *R. triquetrus, R. loreus, R. squarrosus* |
| *Sciuro-hypnum spp.* | *S. reflexum, S. sp* |
| *Sphagnum spp.* | *S. angustifolium, S. capillifolium, S. compactum, S. girgensohnii, S. russowii* |
| *Tetraplodon spp.* | *T. angustatus, T. mnioides* |
| *Hepaticae* | *Anthelia julacea, A. juratzkana; Barbilophozia hatcheri, B. sudetica, B. sp.; Blepharostoma trichophyllum; Calypogeia integristipula, C. neesiana; Cephalozia bicuspidata, C. sp.; Cephaloziella sp.; Diplophyllum albiscans, D. taxifolium; Fuscocephaloziopsis leucantha, F. lunulifolia; Gymnomitrium brevissimum, G. concinnatum, G. corallioides; Hygrobiella laxifolia; Lophozia bicrenata, L. longidens, L. ventricosa, L. wenzelii, L. sp; Marsupella apiculata; Mylia anomala; Nardia geoscyphus, N. scalaris; Neoorthocaulis attenuatus, N. binsteadii, N. floerkei; Odontoschisma francisi, O. elongatum; Orthocaulis atlanticus; Prasanthus suecicus; Scapania sp.; Schljakovia kunzeana; Solenostoma sphaerocarpum; Sphenolobus minutus, S. saxicola; Tetralophozia setiformis; Trilophozia quinquedentate* |
| *Barbilophopzia lycopodioides* | *B. lycopodioides, B. barbata* (most often *B. lycopodioides*) |

| **Lichens** | **Include** |
|---|---|
| *Cladonia arbuscula* | *C. mitis* |
| *Cladonia spp.* | *C. amaurocraea, C. bellidiflora, C. botrytes, C. carneola, C. cenotea, C. chlorophaea, C. coccifera, C. cornuta, C. crispata, C. cyanipes, C. decorticata, C. deformis, C. digitata, C. fimbriata, C. furcata, C. gracilis, C. grayi, C. macilenta, C. macrophyllodes, C. merochlorophaea, C. phyllophora, C. pleurota, C. pyxidata, C. squamosa, C. stricta, C. subfurcata, C. sulphurina, C. trassii, C. verticillata* |
| *Cetraria ssp.* | *C. aculeata, C. ericetorum, C. islandica, C. nigricans, C. odontella; Cetrariella delisei* |
| *Collema ssp.* | *Collema sp.* |
| *Flavocetraria ssp.* | *F. cucullata, F. nivalis* |
| *Hypogymnia ssp.* | *H. bitteri, H. physodes, H. vittata; Parmelia saxatilis, P. sulcata, P. sp.* |
| *Nephroma ssp.* | *N. arcticum, expallidum* |
| *Peltigera ssp.* | *P. apthosa, P. canina, P. degenii, P. didactyla, P. latiloba, P. leucophlebia, P. malacea, P. neckeri, P. neopolydactyla, P. polydactylon, P. rufescens, P. scabrosa, P. sp.* |
| *Sphaerophors spp.* | *S. fragilis, S. globusus* |
| *Stereocaulon spp.* | *S. alpinum, S. arcticum, S. condensatum, S. cumulatum, S. glareosum, S. paschale, S. rivulorum, S. tomentosum, S. vesuvianum, S. sp.* |

**Table S2.** Species' frequencies and mean absolute covers (original survey/resurvey) across all sites. Statistically significant change (after 9999 permutations) in mean cover is indicated as * p > 0.05, ** p > 0.01, *** p > 0.001. Bryophytes and lichens are treated at the genus level, but in the case of one species making up the genus, the full name of the species is given. The taxon nomenclature follows the FinBIF checklist of Finnish species (2024).

| **Vascular plants** | freq. | mean % | | freq. | mean % | | freq. | mean % | | freq. | mean % |
|---|---|---|---|---|---|---|---|---|---|---|---|
| *Achillea millefolium* | 0 / 3 | 0 / + | *Cornus suecica* | 54 / 48 | 1.1 / 0.9 | *Linnaea borealis* | 64 / 69 | 0.3 / 0.2 | *Salix hastata* | 0 / 5 | 0 / +* |
| *Agrostis spp.* | 2 / 7 | + / + | *Dactylorhiza* | | | *Luzula arcuata* | 1 / 0 | + / 0 | *Salix lanata* | 0 / 1 | 0 / + |
| *Alchemilla spp.* | 1 / 1 | + / + | *maculata* | 0 / 1 | + / + | *Luzula multiflora* | 0 / 6 | 0 / +* | *Salix lapponum* | 2 / 5 | + / + |
| *Andromeda polifolia* | 26 / 27 | 0.1 / + | *Deschampsia* | | | *Luzula pilosa* | 1 / 2 | + / + | *Salix myrsinifolia* | 0 / 2 | 0 / + |
| *Antennaria alpina* | 1 / 1 | + / + | *cespitosa* | 8 / 4 | + / + | *Luzula spicata* | 4 / 2 | + / + | *Salix phylicifolia* | 0 / 3 | 0 / + |
| *Antennaria dioica* | 4 / 3 | + / + | *Diapensia lapponica* | 21 / 26 | 0.1 / 0.1 | *Luzula sudetica* | 1 / 0 | + / 0 | *Salix polaris* | 0 / 1 | 0 / + |
| *Anthoxanthum* | | | *Diphasiastrum* | | | *Lycopodium clavatum* | 11 / 20 | + / 0.1 | *Salix reticulata* | 1 / 0 | + / 0 |
| *nipponicum* | 7 / 5 | + / + | *alpinum* | 34 / 35 | 0.1 / 0.1 | *Lysimachia europaea* | 30 / 31 | + / + | *Saussurea alpina* | 5 / 2 | + / + |
| *Anthoxanthum* | | | *Diphasiastrum* | | | *Maianthemum biflora* | 1 / 2 | + / + | *Scorzoneroides* | | |
| *monticola* | 1 / 0 | + / 0 | *complanatum* | 12 / 0 | 0.3 / 0*** | *Melampyrum pratense* | 3 / 9 | + / +* | *autumnalis* | 2 / 1 | + / + |
| *Arctous alpina* | 65 / 87 | 1.9 / 2.3 | *Drosera rotundifolia* | 0 / 1 | + / + | *Nardus stricta* | 6 / 4 | + / + | *Selaginella* | | |
| *Arctostaphylos uva-ursi* | 3 / 7 | + / + | *Dryas octopetala* | 0 / 2 | 0 / + | *Oreojuncus trifidus* | 75 / 93 | 0.2 / 0.2 | *selaginoides* | 3 / 2 | + / + |
| *Astragalus alpinus* | 2 / 3 | + / + | *Empetrum nigrum* | 268 / 273 | 28.4 / 38.3*** | *Orthilia secunda* | 1 / 1 | + / + | *Sibbaldia* | | |
| *Astragalus frigidus* | 0 / 2 | + / + | *Equisetum arvense* | 1 / 2 | + / 0.1 | *Parnassia palustris* | 1 / 0 | + / 0 | *procumbens* | 1 / 2 | + / + |
| *Avenella flexuosa* | 134 / 148 | 0.9 / 0.7 | *Equisetum pratense* | 2 / 2 | + / + | *Pedicularis lapponica* | 56 / 39 | 0.1 / +** | *Silene acaulis* | 1 / 3 | + / + |
| *Bartsia alpina* | 5 / 2 | + / + | *Equisetum sylvaticum* | 1 / 1 | + / + | *Phyllodoce caerulea* | 42 / 59 | 0.5 / 0.6 | *Solidago virgaurea* | 46 / 39 | 0.1 / 0.1 |
| *Betula pubescens* | | | *Erigeron uniflorus* | 0 / 1 | 0 / + | *Picea abies* | 1 / 6 | + / + | *Sorbus aucuparia* | 4 / 5 | + / + |
| *ssp. czerepanovii* | 4 / 30 | + / 0.3*** | *Eriophorum* | | | *Pinguicula alpina* | 2 / 0 | + / 0 | *Spinulum annotinum* | 36 / 24 | 0.1 / 0.1 |
| *Betula nana* | 160 / 157 | 6.7 / 9.8*** | *angustiolium* | 1 / 0 | + / 0 | *Pinguicula vulgaris* | 7 / 3 | + / + | *Taraxacum spp.* | 1 / 0 | + / 0 |
| *Betula pubescens* | 11 / 6 | + / + | *Eriophorum* | | | *Pinus sylvestris* | 2 / 11 | + / +** | *Thalictrum alpinum* | 2 / 1 | + / + |
| *Bistorta vivipara* | 15 / 11 | + / + | *vaginatum* | 2 / 0 | + / + | *Plantago major* | 1 / 0 | + / 0 | *Tofieldia pusilla* | 3 / 3 | + / + |
| *Calamagrostis* | | | *Euphrasia frigida* | 4 / 1 | + / + | *Poa spp.* | 0 / 3 | 0 / + | *Trichophorum* | | |
| *lapponica* | 65 / 43 | 0.3 / 0.1** | *Festuca ovina* | 48 / 50 | 0.2 / 0.1 | *Populus tremula* | 3 / 0 | + / 0 | *cespitosum* | 7 / 5 | + / + |
| *Calluna vulgaris* | 39 / 60 | 4.4 / 4.6 | *Festuca rubra* | 4 / 0 | + / + | *Pyrola minor* | 6 / 0 | + / 0* | *Trollius europaeus* | 3 / 1 | + / + |
| *Campanula* | | | *Festuca vivipara* | 5 / 4 | + / + | *Pyrola rotundifolia* | 0 / 1 | 0 / + | *Vaccinium* | | |
| *rotundifolia* | 3 / 4 | + / + | *Geranium sylvaticum* | 2 / 1 | + / + | *Rhinanthus minor* | 1 / 0 | + / 0 | *myrtillus* | 147 / 175 | 13.5 / 9.1*** |
| *Carex bigelowii* | 65 / 52 | 0.3 / 0.2 | *Gnaphalium supinum* | 2 / 1 | + / + | *Rhodiola rosea* | 0 / 1 | 0 / + | *Vaccinium* | | |
| *Carex brunnescens* | 5 / 5 | + / + | *Gymnocarpium* | | | *Rhododendron* | | | *oxycoccos* | 0 / 1 | 0 / + |
| *Carex globularis* | 5 / 4 | + / + | *dryopteris* | 14 / 4 | + / + | *tomentosum* | 4 / 7 | + / 0.1* | *Vaccinium* | | |
| *Carex nigra* | 1 / 1 | + / + | *Harrimanella hypnoides* | 2 / 5 | + / + | *Rubus chamaemorus* | 22 / 23 | 0.1 / 0.3* | *uliginosum* | 171 / 177 | 2.3 / 4.5*** |
| *Carex paupercula* | 1 / 0 | + / + | *Hieracium spp.* | 16 / 15 | + / + | *Rubus idaeus* | 0 / 1 | 0 / + | *Vaccinium* | | |
| *Carex vaginata* | 7 / 22 | + / +* | *Huperzia selago* | 7 / 12 | + / + | *Rubus saxatilis* | 0 / 1 | 0 / + | *vitis-idaea* | 253 / 253 | 3.3 / 2.4** |
| *Cassiope tetragona* | 6 / 6 | + / 0.1 | *Juncus filiformis* | 1 / 3 | + / + | *Rumex acetosa* | 6 / 1 | + / + | *Vicia cracca* | 0 / 3 | 0 / + |
| *Chamaenerion* | | | *Juniperus communis* | 22 / 39 | 0.1 / 0.6** | *Salix herbacea* | 31 / 27 | 0.3 / 0.2 | *Violabiflora* | 3 / 2 | + / + |
| *angustifolium* | 2 / 5 | + / 0.1 | *Kalmia procumbens* | 47 / 64 | 0.9 / 0.8 | *Salix glauca* | 35 / 26 | 0.1 / 0.1 | *Viola canina* | 0 / 1 | 0 / + |

| **Bryophytes** | freq. | mean % |
|---|---|---|
| Mosses: | | |
| *Abietinella abietina* | 0 / 1 | 0 / + |
| *Andreaea rupestris* | 2 / 4 | + / + |
| *Aulacomnium spp.* | 17 / 9 | 0.3 / +** |
| *Sciuro-hypnum spp.* | 3 / 3 | + / + |
| *Ptychostomum spp.* | 6 / 4 | + / + |
| *Ceratodon purpureus* | 1 / 0 | + / 0 |
| *Climacium dendroides* | 1 / 0 | + / 0 |
| *Conostomum tetragonum* | 2 / 1 | + / + |
| *Cynodontium strumiferum* | 0 / 1 | 0 / + |
| *Dicranella ssp.* | 0 / 1 | 0 / + |
| *Hymenoloma crispulum* | 0 / 3 | 0 / + |
| *Dicranum spp.* | 260 / 265 | 12.7 / 11.7 |
| *Ditrichum spp.* | 1 / 1 | + / + |
| *Hylocomiastrum pyrenaicum* | 2 / 1 | + / + |
| *Hylocomium splendens* | 77 / 79 | 5.2 / 4.5 |
| *Kiaeria spp.* | 8 / 9 | 0.5 / +* |
| *Mnium spp.* | 1 / 2 | + / + |
| *Oligotrichum hercynicum* | 3 / 0 | + / 0 |
| *Oncophorus wahlenbergii* | 1 / 0 | + / 0 |
| *Plagiomnium ellipticum* | 1 / 1 | + / + |
| *Plagiothecium spp.* | 3 / 1 | + / + |
| *Pleurozium schreberi* | 162 / 189 | 7.9 / 12.5*** |
| *Pogonatum spp.* | 7 / 4 | + / + |
| *Pohlia spp.* | 129 / 95 | 0.1 / 0.1 |
| *Polytrichum spp.* | 206 / 197 | 2.3 / 1.1*** |
| *Racomitrium spp.* | 31 / 39 | 0.3 / 0.3 |
| *Rhytidiadelphus spp.* | 12 / 17 | 0.2 / 0.5 |
| *Rhytidium rugosum* | 4 / 6 | + / + |
| *Sanionia uncinata* | 30 / 14 | 0.8 / 0.1* |
| *Sphagnum spp.* | 6 / 7 | 0.2 / 0.1 |
| *Straminergon stramineum* | 1 / 0 | 0.1 / + |
| *Tetraplodon spp.* | 3 / 1 | + / + |

| **Bryophytes** | freq. | mean % |
|---|---|---|
| Liverworts: | | |
| *Barbilophozia lycopodioides* | 142 / 181 | 3.4 / 2.7 |
| *Hepaticae* | 184 / 160 | 2.4 / 0.9*** |
| *Ptidilium ciliare* | 145 / 160 | 3.5 / 4.0 |

| **Lichens** | freq. | mean % |
|---|---|---|
| Reindeer lichens: | | |
| *Cladonia arbuscula* | 232 / 233 | 2.7 / 1.6** |
| *Cladonia rangiferina* | 144 / 205 | 0.3 / 0.7*** |
| *Cladonia stellaris* | 72 / 83 | 0.6 / 0.2** |
| Foliose and fruticose lichens: | | |
| *Cladonia spp.* | 255 / 254 | 2.2 / 2.5 |
| *Alectoria ochroleuca* | 33 / 31 | 0.1 / 0.1 |
| *Bryocaulon divergens* | 17 / 15 | + / + |
| *Cetraria spp.* | 201 / 178 | 0.8 / 0.4*** |
| *Collema spp.* | 2 / 0 | + / 0 |
| *Flavocetraria spp.* | 122 / 112 | 0.7 / 0.6 |
| *Gowardia nigricans* | 30 / 24 | + / + |
| *Hypogymnia spp.* | 23 / 8 | 0.1 / +* |
| *Lobaria linita* | 3 / 4 | + / + |
| *Nephroma spp.* | 98 / 74 | 0.5 / 0.8 |
| *Peltigera* | 113 / 90 | 0.6 / 0.5 |
| *Pseudephebe pubescens* | 14 / 9 | + / + |
| *Solorina crocea* | 19 / 20 | + / + |
| *Sphaerophorus spp.* | 59 / 57 | 0.3 / 0.2 |
| *Stereocaulon spp.* | 134 / 125 | 6.2 / 1.0*** |
| *Thamnolia.vermicularis* | 28 / 26 | + / + |
| Crustaceous lichens | 169 / 136 | 2.3 / 1.1** |

**Reference**: FinBIF (2024): The FinBIF checklist of Finnish species 2023. Finnish Biodiversity Information Facility, Finnish Museum of Natural History, University of Helsinki, Helsinki. https://laji.fi/en/theme/checklist

**Table S3.** Number of different plots with different combinations of biogeographical zone and continentality class. Note that subcontinental northern boreal heathlands are relatively poorly represented in Finland and northern Norway and that *Betula nana* -dominated types are typical to hemioroarctic conditions only and *Calluna vulgaris* -dominated types to northern boreal conditions (Haapasaari, 1988).

| | Northern boreal | | | Hemioroarctic | | | Oroarctic | | |
|---|---|---|---|---|---|---|---|---|---|
| | Subcont. | Indiffer. | Oceanic | Subcont. | Indiffer. | Oceanic | Subcont. | Indiffer. | Oceanic |
| E. nigrum | - | 21 | 9 | 13 | - | 16 | 9 | 19 | 17 |
| V. myrtillus | - | 22 | 12 | - | 4 | 15 | 13 | 13 | 8 |
| B. nana | - | - | - | 7 | 8 | 16 | - | - | - |
| C. vulgaris | - | 23 | 6 | - | - | - | - | - | - |
| Windswept | 2 | - | 1 | 1 | 1 | 3 | 2 | - | 14 |

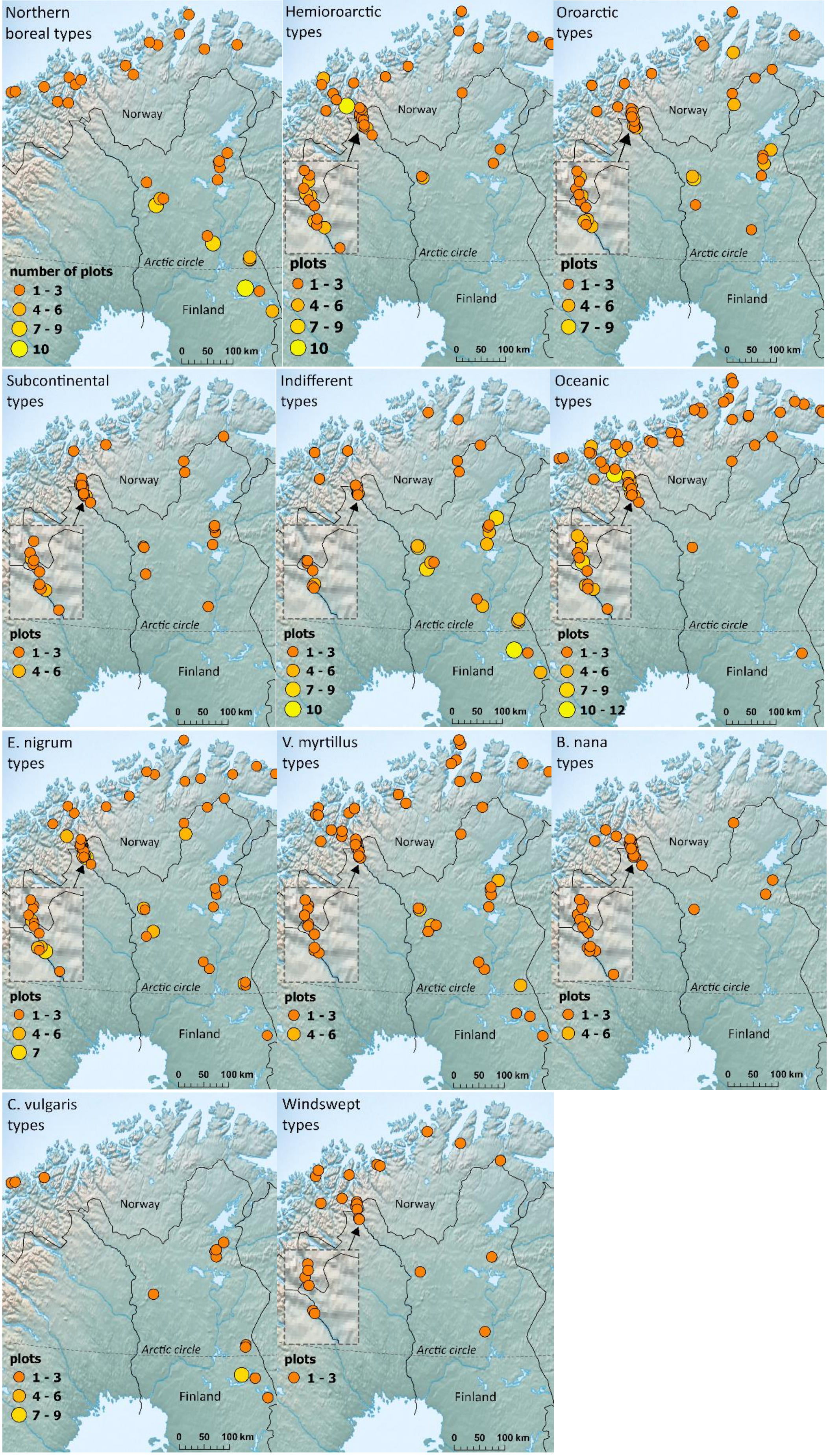


**Figure S1.** Distribution of vegetation plots after classification into different biogeographical zones, continentality classes and habitat types.

**Table S4.** Results from PERMDISP analyses.

| | All species | | | | Vascular plants | | | | Bryophytes | | | | Lichens | | | |
|---|---|---|---|---|---|---|---|---|---|---|---|---|---|---|---|---|
| | Avg. distance to median | | | | Avg. distance to median | | | | Avg. distance to median | | | | Avg. distance to median | | | |
| | old | new | F | *p.perm* | old | new | F | *p.perm* | old | new | F | *p.perm* | old | new | F | *p.perm* |
| All sites ($df_{1, 548}$) | 0.53 | 0.46 | 77.7 | 0.001 | 0.50 | 0.41 | 57.1 | 0.001 | 0.57 | 0.55 | 5.2 | 0.024 | 0.59 | 0.55 | 29.7 | 0.001 |
| Biogeographical zones: | | | | | | | | | | | | | | | | |
| Northern boreal ($df_{1,190}$) | 0.51 | 0.44 | 20.8 | 0.001 | 0.48 | 0.40 | 16.6 | 0.001 | 0.57 | 0.52 | 6.1 | 0.014 | 0.55 | 0.54 | 0.7 | 0.401 |
| Hemioroarctic ($df_{1,166}$) | 0.51 | 0.43 | 26.9 | 0.001 | 0.48 | 0.37 | 28.5 | 0.001 | 0.53 | 0.54 | 0.2 | 0.658 | 0.59 | 0.55 | 11.9 | 0.001 |
| Oroarctic ($df_{1,188}$) | 0.51 | 0.42 | 32.6 | 0.001 | 0.48 | 0.38 | 24.6 | 0.001 | 0.55 | 0.52 | 2.1 | 0.148 | 0.57 | 0.51 | 20.6 | 0.001 |
| Continentality: | | | | | | | | | | | | | | | | |
| Subcontinental ($df_{1,92}$) | 0.44 | 0.45 | 0.1 | 0.948 | 0.47 | 0.41 | 2.7 | 0.103 | 0.48 | 0.51 | 1.1 | 0.288 | 0.43 | 0.51 | 2.6 | 0.111 |
| Indifferent ($df_{1,220}$) | 0.50 | 0.44 | 19.6 | 0.001 | 0.48 | 0.42 | 12.2 | 0.001 | 0.55 | 0.50 | 8.4 | 0.004 | 0.50 | 0.48 | 1.0 | 0.318 |
| Oceanic ($df_{1,232}$) | 0.52 | 0.43 | 30.2 | 0.001 | 0.49 | 0.36 | 40.3 | 0.001 | 0.57 | 0.55 | 1.6 | 0.201 | 0.59 | 0.58 | 0.6 | 0.435 |
| Habitat types: | | | | | | | | | | | | | | | | |
| E. nigrum ($df_{1,206}$) | 0.41 | 0.39 | 1.7 | 0.190 | 0.30 | 0.31 | 0.3 | 0.613 | 0.54 | 0.52 | 1.5 | 0.230 | 0.58 | 0.53 | 12.3 | 0.001 |
| V. myrtillus ($df_{1,172}$) | 0.41 | 0.42 | 0.9 | 0.334 | 0.31 | 0.37 | 10.1 | 0.002 | 0.52 | 0.53 | 0.5 | 0.501 | 0.59 | 0.53 | 11.7 | 0.001 |
| B. nana ($df_{1,60}$) | 0.41 | 0.36 | 5.1 | 0.027 | 0.28 | 0.28 | 0.1 | 0.818 | 0.54 | 0.50 | 1.9 | 0.176 | 0.59 | 0.54 | 7.5 | 0.007 |
| C. vulgaris ($df_{1,56}$) | 0.37 | 0.39 | 1.0 | 0.317 | 0.26 | 0.31 | 2.8 | 0.100 | 0.56 | 0.52 | 1.7 | 0.201 | 0.46 | 0.51 | 1.3 | 0.262 |
| Windswept ($df_{1,46}$) | 0.47 | 0.44 | 1.2 | 0.285 | 0.46 | 0.38 | 2.5 | 0.122 | 0.50 | 0.56 | 2.5 | 0.152 | 0.48 | 0.47 | 0.1 | 0.927 |

**Table S5.** Results from cover (%) variable fits to PcOA (main document Fig. 2a)

| fitted variable | PCoA1 | PCoA2 | $r^2$ | Pr(>r) |
|---|---|---|---|---|
| Empetrum nigrum (%) | -0.85868 | -0.51251 | 0.8651 | 0.001 |
| Betula nana (%) | 0.28101 | -0.95971 | 0.0017 | 0.518 |
| Calluna vulgaris (%) | 0.4894 | 0.87206 | 0.0143 | 0.264 |
| Vaccinium myrtillus (%) | 0.77477 | -0.63224 | 0.7150 | 0.001 |
| Vaccinium vitis-idaea (%) | -0.59827 | -0.8013 | 0.0198 | 0.320 |
| vascular plants (%) | -0.25488 | -0.96697 | 0.5813 | 0.001 |
| bryophytes (%) | 0.19629 | -0.98055 | 0.3452 | 0.001 |
| lichens (%) | 0.22559 | 0.97422 | 0.2604 | 0.001 |

**Table S6.** Changes in species richness (SPR) and evenness (EVE) over time.

| | All species | | | | | | Vascular plants | | | | | |
|---|---|---|---|---|---|---|---|---|---|---|---|---|
| | SPR | | | EVE | | | SPR | | | EVE | | |
| | T1 | T2 | *p.perm* | T1 | T2 | *p.perm* | T1 | T2 | *p.perm* | T1 | T2 | *p.perm* |
| All sites | 20.2 | 20.4 | 0.496 | 0.56 | 0.59 | 0.008 | 8.5 | 9.0 | 0.028 | 0.47 | 0.51 | 0.001 |
| Biogeographical zones: | | | | | | | | | | | | |
| Northern boreal | 18.4 | 17.8 | 0.180 | 0.55 | 0.58 | 0.069 | 8.0 | 8.5 | 0.126 | 0.43 | 0.52 | 0.001 |
| Hemioroarctic | 20.8 | 21.8 | 0.062 | 0.57 | 0.60 | 0.074 | 8.8 | 9.5 | 0.123 | 0.49 | 0.51 | 0.310 |
| Oroarctic | 21.4 | 27.7 | 0.545 | 0.57 | 0.58 | 0.307 | 8.6 | 8.9 | 0.483 | 0.49 | 0.51 | 0.317 |
| Continentality: | | | | | | | | | | | | |
| Subcontinental | 20.9 | 21.9 | 0.223 | 0.60 | 0.61 | 0.479 | 8.0 | 9.0 | 0.118 | 0.51 | 0.53 | 0.582 |
| Indifferent | 19.1 | 18.6 | 0.294 | 0.58 | 0.60 | 0.262 | 7.8 | 8.6 | 0.011 | 0.48 | 0.55 | 0.001 |
| Oceanic | 20.8 | 21.5 | 0.233 | 0.53 | 0.57 | 0.017 | 9.3 | 9.3 | 0.970 | 0.45 | 0.47 | 0.207 |
| Habitat types: | | | | | | | | | | | | |
| E. nigrum | 20.2 | 20.1 | 0.807 | 0.52 | 0.57 | 0.003 | 7.4 | 7.9 | 0.135 | 0.40 | 0.47 | 0.001 |
| V. myrtillus | 19.6 | 20.4 | 0.127 | 0.60 | 0.60 | 0.507 | 9.3 | 10.3 | 0.028 | 0.50 | 0.56 | 0.001 |
| B. nana | 20.2 | 20.7 | 0.670 | 0.59 | 0.61 | 0.511 | 9.4 | 9.1 | 0.774 | 0.52 | 0.54 | 0.541 |
| C. vulgaris | 20.3 | 18.0 | 0.016 | 0.58 | 0.59 | 0.754 | 9.5 | 9.0 | 0.508 | 0.46 | 0.53 | 0.031 |
| Windswept | 21.7 | 24.2 | 0.043 | 0.58 | 0.58 | 0.848 | 7.6 | 8.6 | 0.162 | 0.62 | 0.46 | 0.005 |

| | Bryophytes | | | | | | Lichens | | | | | |
|---|---|---|---|---|---|---|---|---|---|---|---|---|
| | SPR | | | EVE | | | SPR | | | EVE | | |
| | T1 | T2 | *p.perm* | T1 | T2 | *p.perm* | T1 | T2 | *p.perm* | T1 | T2 | *p.perm* |
| All sites | 5.3 | 5.3 | 0.943 | 0.52 | 0.50 | 0.286 | 6.4 | 6.1 | 0.068 | 0.6 | 0.69 | 0.001 |
| Biogeographical zones: | | | | | | | | | | | | |
| Northern boreal | 5.0 | 4.9 | 0.646 | 0.51 | 0.48 | 0.509 | 5.5 | 4.4 | 0.001 | 0.63 | 0.66 | 0.303 |
| Hemioroarctic | 5.8 | 5.8 | 0.947 | 0.55 | 0.52 | 0.353 | 6.2 | 6.5 | 0.281 | 0.57 | 0.71 | 0.002 |
| Oroarctic | 5.1 | 5.2 | 0.624 | 0.50 | 0.49 | 0.751 | 7.6 | 7.6 | 0.900 | 0.61 | 0.71 | 0.001 |
| Continentality: | | | | | | | | | | | | |
| Subcontinental | 4.8 | 4.9 | 0.861 | 0.57 | 0.53 | 0.334 | 8.2 | 8.0 | 0.669 | 0.45 | 0.69 | 0.001 |
| Indifferent | 4.8 | 4.6 | 0.254 | 0.47 | 0.49 | 0.673 | 6.5 | 5.5 | 0.001 | 0.66 | 0.73 | 0.001 |
| Oceanic | 5.9 | 6.2 | 0.334 | 0.54 | 0.50 | 0.073 | 5.6 | 6.0 | 0.165 | 0.61 | 0.66 | 0.238 |
| Habitat types: | | | | | | | | | | | | |
| E. nigrum | 5.3 | 5.2 | 0.428 | 0.53 | 0.50 | 0.397 | 7.5 | 7.1 | 0.133 | 0.60 | 0.72 | 0.001 |
| V. myrtillus | 5.2 | 5.3 | 0.958 | 0.52 | 0.46 | 0.065 | 5.1 | 4.9 | 0.435 | 0.60 | 0.67 | 0.087 |
| B. nana | 5.5 | 5.5 | 0.999 | 0.54 | 0.51 | 0.540 | 5.3 | 6.0 | 0.117 | 0.64 | 0.73 | 0.182 |
| C. vulgaris | 5.2 | 5.0 | 0.707 | 0.49 | 0.57 | 0.216 | 5.6 | 3.9 | 0.001 | 0.62 | 0.68 | 0.389 |
| Windswept | 5.0 | 6.0 | 0.072 | 0.46 | 0.51 | 0.371 | 9.0 | 9.6 | 0.227 | 0.60 | 0.62 | 0.633 |

**Table S7.** Species' frequencies and mean absolute covers (original survey/resurvey) within different biogeographical zones. Only species with statistically significant change (9999 permutations, p.perm) in cover are listed. * Significant (p > 0.05) after the Holm correction.

| | **N. boreal** (n plots = 96, n species = 126) | | | | | **Hemioroarctic** (n plots = 84, n species = 127) | | | | | **Oroarctic** (n plots = 95, n species = 127) | | | | |
|---|---|---|---|---|---|---|---|---|---|---|---|---|---|---|---|
| species | freq. T1 / T2 | cover (%) T1 / T2 | Δ cover | *t* | p.perm | freq. T1 / T2 | cover (%) T1 / T2 | Δ cover | *t* | p.perm | freq. T1 / T2 | cover (%) T1 / T2 | Δ cover | *t* | p.perm |
| *Betula pub. ssp. czerepanovii* | 2 / 13 | + / 0.5 | +0.5 | 1.9 | 0.002 | 0 / 9 | 0 / 0.2 | +0.2 | 2.1 | 0.002 | | | | | |
| *Betula nana* | | | | | | | | | | | 78 / 74 | 3.2 / 9.6 | +6.4 | 4.9 | 0.001* |
| *Calamagrostis lapponica* | | | | | | 33 / 20 | 0.3 / 0.1 | -0.2 | -2.5 | 0.004 | | | | | |
| *Calluna vulgaris* | | | | | | | | | | | 5 / 11 | 0.1 / 0.8 | +0.7 | 1.8 | 0.002 |
| *Cornus suecica* | | | | | | | | | | | 1 / 8 | + / 0.4 | +0.4 | 1.1 | 0.008 |
| *Diphasiastrum complanatum* | 10 / 0 | 0.1 / 0 | -0.1 | -1.4 | 0.002 | | | | | | | | | | |
| *Empetrum nigrum* | 95 / 95 | 28.7 / 42.2 | +13.5 | 4.7 | 0.001* | 79 / 83 | 28.9 / 38.9 | +10.0 | 3.1 | 0.001 | 94 / 95 | 27.5 / 33.7 | +6.2 | 2.7 | 0.004 |
| *Juniperus communis* | 11 / 19 | + / 1.4 | +1.4 | 2.2 | 0.001 | | | | | | | | | | |
| *Oreojuncus trifidus* | 12 / 16 | + / 0.2 | +0.2 | 2.1 | 0.008 | | | | | | | | | | |
| *Pedicularis lapponica* | | | | | | 26 / 21 | 0.1 / + | -0.1 | -2.3 | 0.007 | | | | | |
| *Pinus sylvestris* | 2 / 9 | + / 0.1 | +0.1 | 1.8 | 0.002 | | | | | | | | | | |
| *Vaccinium myrtillus* | | | | | | 36 / 48 | 12.4 / 6.7 | -5.7 | -2.5 | 0.007 | 38 / 50 | 14.4 / 8.6 | -5.7 | -3.9 | 0.001* |
| *Vaccinium uliginosum* | 57 / 62 | 1.8 / 6.3 | +4.5 | 4.9 | 0.001* | | | | | | | | | | |
| *Vaccinium vitis-idaea* | | | | | | 79 / 79 | 3.5 / 2.3 | -1.2 | -2.5 | 0.007 | 85 / 81 | 2.6 / 1.4 | -1.2 | -3.6 | 0.001* |
| *Hepaticae* | | | | | | | | | | | 75 / 64 | 4.2 / 1.4 | -2.8 | -2.5 | 0.001 |
| *Pleurozium schreberi* | 70 / 84 | 9.9 / 18.3 | +8.5 | 3.6 | 0.001* | | | | | | 30 / 40 | 0.3 / 2.3 | +1.9 | 2.8 | 0.002 |
| *Polytrichum spp.* | 63 / 48 | 0.9 / 0.3 | -0.6 | -2.4 | 0.001 | | | | | | 76 / 83 | 4.4 / 1.6 | -2.8 | -2.7 | 0.001 |
| *Sanionia uncinata* | 12 / 5 | 0.7 / + | -0.7 | -1.3 | 0.006 | | | | | | | | | | |
| *Cetraria spp.* | 74 / 56 | 0.8 / 0.3 | -0.5 | -2.6 | 0.004 | | | | | | 78 / 78 | 1.1 / 0.6 | -0.5 | -2.7 | 0.003 |
| *Cladonia arbuscula* | 78 / 74 | 4.4 / 1.5 | -2.9 | -3.8 | 0.001* | | | | | | | | | | |
| *Cladonia rangiferina* | 52 / 74 | 0.3 / 0.8 | +0.5 | 3.4 | 0.001* | | | | | | | | | | |
| *Cladonia stellaris* | 40 / 27 | 0.7 / 0.1 | -0.6 | -2.3 | 0.001 | | | | | | | | | | |
| *Flavocetraria spp.* | | | | | | 40 / 35 | 0.2 / 0.7 | +0.5 | 2.6 | 0.003 | | | | | |
| *Peltigera spp.* | | | | | | | | | | | 33 / 28 | 0.7 / 0.3 | -0.4 | -1.9 | 0.009 |
| *Stereocaulon spp.* | | | | | | 47 / 54 | 12.8 / 1.7 | -11.1 | -4.5 | 0.001* | 56 / 54 | 5.8 / 1.2 | -4.6 | -3.3 | 0.001 |
| *crustaceous lichens* | | | | | | | | | | | 80 / 73 | 5.0 / 2.0 | -3.0 | -3.1 | 0.001* |

**Table S8.** Species' frequencies and mean absolute covers (original survey/resurvey) within different continentality/humidity classes. Only species with statistically significant change (9999 permutations, p.perm) in cover are listed. * Significant (p > 0.05) after the Holm correction.

| | **Subcontinental/arid** (n plots = 47, n species = 99) | | | | | **Indifferent** (n plots = 111, n species = 108) | | | | | **Oceanic/humid** (n plots = 117, n species = 152) | | | | |
|---|---|---|---|---|---|---|---|---|---|---|---|---|---|---|---|
| species | freq. T1 / T2 | cover (%) T1 / T2 | Δ cover | *t* | p.perm | freq. T1 / T2 | cover (%) T1 / T2 | Δ cover | *t* | p.perm | freq. T1 / T2 | cover (%) T1 / T2 | Δ cover | *t* | p.perm |
| *Betula pub. ssp. czerepanovii* | | | | | | | | | | | 1 / 20 | + / 0.3 | +0.3 | 2.5 | 0.001* |
| *Betula nana* | 36 / 37 | 7.1 / 12.2 | +5.1 | 1.6 | 0.004 | | | | | | 75 / 72 | 8.6 / 12.5 | +3.9 | 3.2 | 0.001 |
| *Diphasiastrum complanatum* | | | | | | 10 / 0 | 0.1 / 0 | -0.1 | -1.4 | 0.002 | | | | | |
| *Empetrum nigrum* | 45 / 46 | 17.7 / 29.3 | +11.6 | 4.0 | 0.001* | 110 / 110 | 23.9 / 33.6 | +9.7 | 4.3 | 0.001* | 113 / 117 | 36.8 / 46.3 | +9.5 | 3.2 | 0.001 |
| *Oreojuncus trifidus* | | | | | | 29 / 35 | 0.1 / 0.3 | +0.2 | 2.4 | 0.007 | | | | | |
| *Pedicularis lapponica* | | | | | | | | | | | 32 / 24 | 0.1 / + | -0.1 | -2.4 | 0.005 |
| *Pinus sylvestris* | | | | | | 1 / 10 | + / 0.1 | +0.1 | 1.7 | 0.002 | | | | | |
| *Vaccinium myrtillus* | | | | | | | | | | | 52 / 64 | 13.7 / 5.3 | -8.4 | -4.6 | 0.001* |
| *Vaccinium uliginosum* | | | | | | 59 / 67 | 1.9 / 4.8 | +2.9 | 3.7 | 0.001* | 89 / 86 | 3.1 / 5.2 | +2.1 | 2.6 | 0.005 |
| *Vaccinium vitis-idaea* | 45 / 43 | 3.1 / 1.6 | -1.5 | -2.9 | 0.001 | | | | | | | | | | |
| *Aulacomnium palustre* | | | | | | | | | | | 14 / 9 | 0.7 / + | -0.7 | -1.5 | 0.008 |
| *Hepaticae* | | | | | | | | | | | 79 / 82 | 4.8 / 1.6 | -3.2 | -3.1 | 0.001* |
| *Pleurozium schreberi* | | | | | | | | | | | 78 / 93 | 9.2 / 16.9 | +7.8 | 3.4 | 0.001* |
| *Polytrichum spp.* | | | | | | 80 / 72 | 2.6 / 0.7 | -1.9 | -2.9 | 0.001* | | | | | |
| *Cetraria spp.* | | | | | | 96 / 89 | 1.1 / 0.6 | -0.5 | -2.7 | 0.003 | | | | | |
| *Cladonia arbuscula* | | | | | | 107 / 96 | 5.2 / 1.8 | -3.4 | -4.6 | 0.001* | | | | | |
| *Cladonia rangiferina* | | | | | | | | | | | 40 / 76 | 0.1 / 0.7 | +0.6 | 3.9 | 0.001* |
| *Cladonia stellaris* | | | | | | 54 / 48 | 0.9 / 0.3 | -0.6 | -2.3 | 0.006 | 1 / 16 | + / 0.1 | +0.1 | 2.8 | 0.001* |
| *Cladonia spp.* | | | | | | | | | | | 100 / 101 | 0.7 / 1.5 | +0.8 | 2.2 | 0.006 |
| *Nephroma spp.* | | | | | | | | | | | 48 / 48 | 0.4 / 1.2 | +0.8 | 2.5 | 0.004 |
| *Peltigera spp.* | 16 / 19 | 0.1 / 0.6 | +0.5 | 2.6 | 0.006 | 36 / 13 | 0.4 / 0.1 | -0.3 | -2.0 | 0.002 | | | | | |
| *Stereocaulon spp.* | 38 / 36 | 30.4 / 2.1 | -28.3 | -7.8 | 0.001* | 49 / 32 | 2.1 / 0.3 | -1.9 | -3.0 | 0.001* | 47 / 57 | 0.4 / 1.3 | +0.9 | 2.5 | 0.003 |
| *crustaceous lichens* | | | | | | | | | | | 52 / 52 | 3.5 / 1.3 | -2.2 | -2.6 | 0.004 |

**Table S9.** Species' frequencies and mean absolute covers (original survey/resurvey) within different habitat types. Only species with statistically significant change (9999 permutations, p.perm) in cover are listed. In *Betula nana* -dominated habitats (n plots = 31, n species = 94), only the cover of *Stereocaulon spp.* changed over time (freq. T1 / T2 = 16 / 20, cover T1 / T2 = 10.2 / 0.8, p.perm = 0.009). * Significant (p > 0.05) after the Holm correction.

| Habitat type: | **E. nigrum** (n plots = 104, n species = 122) | | | | | **V. myrtillus** (n plots = 87, n species = 129) | | | | |
|---|---|---|---|---|---|---|---|---|---|---|
| species | freq. T1 / T2 | cover (%) T1 / T2 | Δ cover | *t* | p.perm | freq. T1 / T2 | cover (%) T1 / T2 | Δ cover | *t* | p.perm |
| *Arctous alpina* | | | | | | 4 / 17 | 0.1 / 2.3 | +2.2 | 2.6 | 0.001* |
| *Avenella flexuosa* | 19 / 40 | 0.1 / 0.3 | +0.2 | 3.1 | 0.001 | 82 / 70 | 2.3 / 1.3 | -0.9 | -3.1 | 0.002 |
| *Betula pubescens ssp. czerepanovii* | 0 / 9 | 0 / 0.4 | +0.4 | 1.3 | 0.002 | 3 / 14 | + / 0.2 | +0.2 | 2.6 | 0.001 |
| *Betula nana* | 68 / 58 | 3.4 / 7.6 | +4.2 | 4 | 0.001* | 36 / 42 | 0.9 / 4.7 | +3.8 | 3.5 | 0.001* |
| *Calluna vulgaris* | 4 / 11 | + / 0.9 | +0.9 | 2.1 | 0.002 | 5 / 16 | 0.2 / 1.8 | +1.6 | 2.4 | 0.003 |
| *Empetrum nigrum* | | | | | | 84 / 87 | 16.2 / 35.0 | +18.7 | 7.3 | 0.001* |
| *Vaccinium myrtillus* | 24 / 48 | 0.2 / 3.6 | +3.4 | 4.7 | 0.001* | 87 / 84 | 39.9 / 21.1 | -18.9 | -8.1 | 0.001* |
| *Vaccinium uliginosum* | | | | | | 48 / 55 | 2.3 / 6.0 | +3.6 | 4.0 | 0.001* |
| *Aulacomnium spp.* | 11 / 6 | 0.8 / + | -0.8 | -1.5 | 0.007 | | | | | |
| *Dicranum spp.* | | | | | | 87 / 87 | 19.9 / 13.3 | -6.6 | -3.1 | 0.002 |
| *Hepaticae* | | | | | | | | | | |
| *Pleurozium schreberi* | 53 / 62 | 4.2 / 8.7 | +4.5 | 3.1 | 0.001 | | | | | |
| *Polytrichum spp.* | 74 / 85 | 2.6 / 1.1 | -1.5 | -2.5 | 0.004 | | | | | |
| *Cetraria spp.* | | | | | | 58 / 57 | 0.7 / 0.3 | -0.4 | -2.4 | 0.006 |
| *Cladonia arbuscula* | | | | | | | | | | |
| *Cladonia rangiferina* | 60 / 81 | 0.2 / 0.8 | +0.6 | 3.5 | 0.001* | | | | | |
| *Peltigera spp.* | 54 / 42 | 1.0 / 0.6 | -0.4 | -2.4 | 0.007 | | | | | |
| *Stereocaulon spp.* | 54 / 50 | 7.1 / 1.3 | -5.9 | -3.4 | 0.001* | 42 / 38 | 6.9 / 0.9 | -6.0 | -3.9 | 0.001* |
| *crustaceous lichens* | | | | | | | | | | |

| Habitat type: | **C. vulgaris** (n plots = 29, n species = 84) | | | | | **Windswept** (n plots = 24, n species = 89) | | | | |
|---|---|---|---|---|---|---|---|---|---|---|
| species | freq. T1 / T2 | cover (%) T1 / T2 | Δ cover | *t* | p.perm | freq. T1 / T2 | cover (%) T1 / T2 | Δ cover | *t* | p.perm |
| *Arctous alpina* | | | | | | | | | | |
| *Avenella flexuosa* | 14 / 20 | 0.1 / 0.6 | +0.5 | 3 | 0.001* | | | | | |
| *Betula pubescens ssp. czerepanovii* | | | | | | | | | | |
| *Betula nana* | | | | | | 18 / 15 | 1.3 / 7.7 | +6.4 | 2.7 | 0.002 |
| *Calluna vulgaris* | | | | | | | | | | |
| *Empetrum nigrum* | 29 / 29 | 14.6 / 32.2 | +17.6 | 4.4 | 0.001* | 23 / 24 | 5.2 / 24.3 | +19.1 | 5.3 | 0.001* |
| *Vaccinium myrtillus* | | | | | | | | | | |
| *Vaccinium uliginosum* | 22 / 19 | 1.6 / 4.3 | +2.7 | 2.1 | 0.008 | | | | | |
| *Aulacomnium spp.* | | | | | | | | | | |
| *Dicranum spp.* | | | | | | | | | | |
| *Hepaticae* | | | | | | 21 / 21 | 17.1 / 6.4 | -10.7 | -2.9 | 0.001* |
| *Pleurozium schreberi* | 22 / 28 | 7.2 / 21.3 | +14.1 | 3.9 | 0.001* | | | | | |
| *Polytrichum spp.* | | | | | | 23 / 23 | 2.6 / 0.5 | -2.1 | -1.7 | 0.001 |
| *Cetraria spp.* | | | | | | 22 / 22 | 1.9 / 0.6 | -1.3 | -2.2 | 0.009 |
| *Cladonia arbuscula* | 29 / 22 | 7.8 / 1.6 | -6.3 | -3.1 | 0.002 | | | | | |
| *Cladonia rangiferina* | 17 / 24 | 0.2 / 0.8 | +0.6 | 2.5 | 0.004 | | | | | |
| *Peltigera spp.* | | | | | | | | | | |
| *Stereocaulon spp.* | 9 / 1 | 0.5 / + | -0.5 | -1.9 | 0.002 | | | | | |
| *crustaceous lichens* | | | | | | 24 / 23 | 16.0 / 5.5 | -10.5 | -3 | 0.002 |
